# Experimental field trials model how the climate crisis will alter the phyllosphere and carposphere fungal communities of *Vitis* sp. L’Acadie Blanc

**DOI:** 10.1101/2024.10.25.620292

**Authors:** Andrew J.C. Blakney, Odile Carisse, Hervé Van Der Heyden, Frederic E. Pitre, Karine Pedneault

## Abstract

The climate crisis is changing temperature regimes worldwide, threatening global viticulture and wine production, as temperature is a primary driver of grape development. In Atlantic Canada, temperatures are projected to increase, inducing premature grape ripening, which can impact their biochemical profiles and, consequently, the quality of the vines and wines produced. Temperature is also a key factor in determining the composition and structure of resident fungal communities on the leaves (phyllosphere) and fruits (carposphere) of grape vines. Therefore, to better understand how these communities might change under potential future temperature regimes, we experimentally manipulated grapevines (*Vitis* sp. cv. L’Acadie blanc) in the field. We used on-the-row mini-greenhouses to increase the temperature at different developmental, or phenological, stages of the fruits, and across the whole season. Phyllosphere and carposphere were sampled at four developmental stages, their DNA was extracted, and the fungal communities were identified via ITS metabarcoding. We found that phyllosphere and carposphere had significantly different community composition, which remained relatively stable throughout plant development. Increased temperature treatments had the most significant effect on fungal phyllosphere communities; we observed that phyllosphere samples exposed to higher temperatures before the onset of ripening maintained more diverse fungal communities throughout development. Our analysis showed that the increase in fungal diversity among phyllopshere communities corresponds to enrichments in potential phytopathogenic fungal taxa. However, this increase in phyllosphere fungal diversity was not conserved at other growth stages when the leaves developed at higher temperatures for the whole season. The results of this study will contribute to better understanding the impact of the climate crisis on grapevine phyllosphere and carposphere fungal community composition and assembly. This will allow producers to better adapt to climate variability and to better understand the role that these communities could play on grapevine health.

## Introduction

The Intergovernmental Panel on Climate Change (IPCC) conservatively estimates global temperatures will increase by 1.4–4.4°C by 2100 depending on greenhouse gas emissions (Masson-Delmotte *et al*., 2021). In Atlantic Canada, temperatures are predicted to increase from 2 to 4°C in the summer and 1.5 to 6°C during the winter (Vasseur & Catto, 2007), resulting in longer, hotter growing seasons, and shorter, warmer, damper winters. These temperature changes will impact agriculture by decreasing yields through heat stress, drought, frequent heavy rainfalls, and the increased populations of phytopathogens (Jobin Poirier *et al*., 2020; Lesk *et al*., 2022; Singh *et al*., 2023).

These climate-induced temperature changes also threaten global viticulture and wine production, as temperature is a primary driver of grape development (Jobin Poirier *et al*., 2020; Ausseil *et al*., 2021). In Atlantic Canada, these increased temperatures could affect grapevine phenology (i.e. developmental stages) and grape ripening, increase disease pressure and, ultimately, affect fruit quality. Grape varieties grown in the region are interspecific crosses between *Vitis vinifera* and *Vitis* species indigenous to North America (e.g. *V*. *labrusca*, *V*. *riparia*), which have been selected for strong cold resistance due to a unique biochemical profile, and an early maturity (Pedneault *et al*. 2013). Typically in the region, berry fruit-set occurs in late June, with growth continuing until the end of August, and the onset of veraison, the ripening stage. Berries will continue to ripen until harvested in mid to late October (Campos-Arguedas *et al*., 2022). During the entirety of berry development, including the berry growth phase, temperature affects the development of flavor, aroma, sugar, and alcohol content of berries (van Leeuwen & Seguin, 2006; Campos-Arguedas *et al*., 2022). For example, early development of berries is characterised by accumulating malic and tartaric acids, and proanthocyanidins. In late development (i.e. ripening) malic acid breaks down, while sugars and various secondary metabolites accumulate (Bonada *et al*., 2015; Campos-Arguedas *et al*., 2022). Our previous work has further experimentally confirmed in field experiments that increasing the temperature during early berry development changes their biochemistry at harvest (Campos-Arguedas *et al*., 2022). Unexpected changes to the biochemical profile of the grapes due to climate-induced temperature variation may thus impact the quality of the grapes and wine.

Temperature, a key factor in the “terroir”, also helps determine the composition and structure of resident microbial communities (van Leeuwan & Seguin, 2006; Malik *et al*., 2019; Jannson & Hofmockel, 2020; Koskella, 2020; Zhu *et al*., 2021; Perreault & Laforest-Lapointe, 2022; Tiedje *et al*., 2022; Di Paola *et al*., 2023). However, as the climate crisis creates more variable temperatures across geographic regions, it also allows new phytopathogens to emerge, or spread into new areas (Malik *et al*., 2019; Jannson & Hofmockel, 2020; Koskella, 2020; Zhu *et al*., 2021; Tiedje *et al*., 2022; Singh *et al*., 2023). Studies have broadly illustrated that bacterial and fungal diversity tracks plant diversity (Leff *et al*., 2018; Nottingham *et al*., 2018), which typically follows global abiotic patterns of temperature and precipitation (Malik *et al*., 2019; He *et al*., 2020; Koskella, 2020; Zhu *et al*., 2021; Tiedje *et al*., 2022). However, fungi appear partial to cooler temperatures, as they make up more biomass of soils at higher latitudes (He *et al*., 2020), but have shown resilience in quickly adapting to warming soils (Romero-Olivares *et al*., 2017). Although soils may be the reservoir for many microbes in the aerial portions of plants (phyllosphere; hereafter used to specify the leaves), important differences can still accrue between above and below-ground communities over the development of the plant host (Jannson & Hofmockel, 2020; Koskella, 2020; Blakney *et al*., 2024b). In general, microbes that colonize the phyllosphere survive under difficult conditions of limited nutrients, exposure to UV radiation and fluctuations in pH, humidity and temperature (Lindow & Brandl, 2003; Koskella, 2020). As such, experimentally modelling how fungal communities of grapevine phyllosphere and fruits (carposphere) change in response to increased temperatures over different phenologies will be useful in understanding the potential impact of the climate crisis on the fungal, or mycobiome, components on the terroir and potential agroecological roles, such as pathogenicity and biological control.

In Canada, there have been very few experiments exploring the relationship between grapevines and their microbial communities, and to our knowledge, none on the hybrid grapevine cultivars grown in Atlantic Canada. Here, we report a field experiment used to test whether changes in temperature patterns at different stages of fruit development (pre-veraison and post-veraison) or a global rise in temperature (whole season) altered the fungal communities of grapevine phyllosphere and carposphere. Our hypothesis was that increased heat treatments would significantly increase fungal diversity of the phyllosphere and carposphere regardless of development stage. As such, we predicted that i) fungal diversity would increase over the growing season in untreated controls, ii) diversity would be higher among heat treated samples than the untreated controls, and iii) diversity would not be different between treated samples from different growth stages. We used on-the-row mini-greenhouses to increase the temperature at different fruit developmental stages of *Vitis* sp. cv. L’Acadie blanc; pre-veraison, post-veraison, or throughout the growing season. Leaves and fruits were sampled at four developmental stages, and fungi were identified via ITS metabarcoding. We measured changes to fungal community composition, and α- and β-diversity across different developmental stages to assess the impact of the increased temperature regimes to the grapevine phyllosphere and carposphere communities. These results will allow us to evaluate the adaptability of the fungal community in the face of increasing temperatures in Atlantic Canada.

## Materials and methods

### 1. Site and experimental design

The field experiment, previously described in Campos-Arguedas *et al*., 2022, was conducted at a commercial vineyard in the Gaspereau Valley, in Wolfville, Nova Scotia (45°4′19″N, 64°17′44″W) during the 2020 growing season. The experimental site was planted with an 11-year-old *Vitis* sp. cv. L’Acadie blanc (Cascade X Seyve-Villard 14-287). The experimental design was a randomized split-plot replicated in five complete blocks. Within each block, plots were split into four treatments, where polycarbonate mini-greenhouses were installed at specific growth stages to simulate increased temperatures, while the control treatment lacked greenhouses throughout the season. The specific growth stages were determined by the modified Eichhorn-Lorenz (EL) system (Coombe, 1995). The whole season heat treatment maintained greenhouses from June 3^rd^ (developmental stage EL-15) to October 9^th^ (harvest, EL-38), which increased growing degree (GDD) days by 197 relative to the control (as previously illustrated by Campos-Arguedas *et al*., 2022); the pre-veraison heat treatment greenhouses were installed on June 3^rd^ until September 10^th^ (onset of veraison, EL-35), with an associated increase of 128 GDD, while the post-veraison heat treatment had greenhouses installed on September 11^th^ until October 9^th^ (harvest, EL-38), with an increase of 44 GDD (see Campos-Arguedas *et al*., 2022 for further design details). These periods were selected based on the double sigmoid curve grape berries follow during their development.

### 2. Crop management and sampling

Grapevines were grown and maintained according to standard management practices, as previously described by Campos-Arguedas *et al* (2022). *Vitis* sp. leaves and berries were sampled at the following growth stages; EL-32 (bunch closure), EL-36 (intermediate fruit sugar levels), EL-37 (fruit not quite ripe), and EL-38 (ripe and harvest). At each of these growth stages, 6-8 berry clusters, and their adjacent leaves, were randomly picked from all four vines in each treatment per block. Sampled material was divided into leaves and fruits and pooled, generating 160 samples (2 compartments (i.e. phyllosphere and carposphere) *4 growth stages *4 treatments *5 replicates). Samples were immediately frozen with liquid nitrogen, transported in dry ice, and stored at -80°C before being shipped to Université de Montréal’s Biodiversity Centre (Montréal, QC, Canada) on dry ice for further processing (Delavaux *et al*., 2020; Blakney *et al*., 2022). As is typical for field experiments, we accounted for the use of the various agricultural management practices (i.e. experimental treatments, fertilizers, pesticides, and management) in the downstream amplicon data by considering each sample and its management history as a unit: i.e. the detection of a given fungal species is considered to be due to the total effect of the management and experimental treatments of the samples.

### 3. DNA extraction from *Vitis* sp. leaf and fruit samples

Total DNA was extracted from leaves and fruits after being ground separately in liquid nitrogen via sterile mortar and pestles. DNeasy Plant DNA Extraction Kits (Qiagen, Germany) were used following the manufacturer’s instructions with ∼160 mg of leaf material and ∼225 mg of fruit material (Lay *et al*., 2018; Blakney *et al*., 2022). No-template extraction negative controls were included with each kit used, to assess the influence of the extraction kits on our sequencing results, and the efficacy of our lab preparation. All extracted DNA samples were qualitatively evaluated by mixing ∼2 μL of each sample with 1 μL of loading dye containing Gel Red (Biotium) and running it on a 0.7 % agarose gel for 50 minutes at 110 V. The no-template extraction negative controls were confirmed not to contain DNA after extraction.

### 4. Amplicon generation and sequencing to estimate fungal communities

To estimate the composition of the fungal communities in the phyllosphere and carposphere across the *Vitis* sp. growth stages, extracted DNA from all samples were used to prepare ITS amplicon libraries, following Illumina’s MiSeq protocols (Bell *et al*., 2016; Lay *et al*., 2018; Blakney *et al*., 2022). First, all DNA samples were diluted 1:10 into 96-well plates. To assess potential bias caused by lab manipulations, sequencing and downstream bioinformatic processing, we also included the no-template extraction control samples.

The prepared plates of the *Vitis* sp. leaf and fruit DNA samples were submitted to Génome Québec (Montréal, Québec) for ITS amplicon generation and sequencing (Bell *et al*., 2016; Lay *et al*., 2018; Blakney *et al*., 2023). Diluted DNA samples were used as templates for PCR amplification with the ITS3-KYO2 forward and ITS4-KYO3 reverse primers, which generated a 430 bp fragment from the ITS2 region, between the 5.8S and LSU regions (Toju *et al*., 2012; Morvan *et al*., 2020). Amplicons were then prepared for paired-end sequencing using Illumina’s MiSeq platform (Génome Québec, Montréal) (Bell *et al*., 2016; Lay *et al*., 2018; Blakney *et al*., 2022). We estimated this should provide a mean of 40 000 ITS reads per sample, which is in line with previous studies that describe bacterial communities (Bell *et al*., 2016; Lay *et al*., 2018; Blakney *et al*., 2022; Morvan *et al*., 2020). Sequencing data and metadata are available at NCBI Bioproject under accession number: PRA1046454.

### 5. Estimating ASV’s from amplicon sequencing

The amplicons generated by Illumina MiSeq were used to estimate the diversity and composition of the fungal communities present in the leaves and fruits of *Vitis* sp. at different growth stages. The integrity and totality of the MiSeq data downloaded from Génome Québec was confirmed using their MD5 checksum protocol (Roy *et al*., 2018). Subsequently, all data was managed, and analyzed in R (4.0.3 R Core Team, 2020), and plotted using ggplot2 (Wickham, 2016).

The raw ITS reads were processed to retain the highest quality reads before ASV inference and taxonomic assignment. Due to the variable length of the ITS region, we first used cutadapt (Martin, 2011) to carefully remove primer sequences from all 1 616 465 raw ITS reads generated from the control samples, and the *Vitis* sp. samples, including any primer sequences generated due to read-through (Blakney *et al*., 2023). The filtered and trimmed reads for the ITS region were then processed through DADA2 for ASV inference (Fig. S1). The default settings were kept throughout the pipeline, except the `dada` inference function, which used the pool =’pseudo’ argument, to increase the likelihood of identifying rare taxa. Consequently, the chimera removal function removeBimeraDenovo included the method =’pooled’ argument (Callahan *et al*., 2016b).

Fungal amplicon sequence variants (ASVs) were assigned taxonomy following the default DADA2 pipeline (Callahan *et al*., 2016b), using the UNITE fungal database (Abarenkov *et al*., 2022), as well as the UNITE database for all eukaryotes (Abarenkov *et al*., 2020), in order to confirm fungal identities (Tedersoo *et al*., 2018). The quality of the data was assessed using the included controls, and any off-target eukaryotic ASVs identified in the fungal data were removed. Rarefaction curves confirmed that we obtained sufficient coverage of the fungi present in both the phyllosphere and carposphere (Fig. S2; Blakney *et al*., 2024a).

### 6. α-diversity of the *Vitis* sp. fungal communities in the phyllosphere and carposphere

First, to visualise the taxonomic diversity, ASVs from the phyllosphere and carposphere were plotted separately as taxa cluster maps using the `heat_tree` function from the `metadcoder` package (Foster *et al*., 2017), where nodes represent phyla to genera: node colours represent the abundance of 16S rRNA reads, while node size indicates the number of unique taxa. Taxa cluster maps facilitate visualizing abundance, as well as diversity across taxonomic hierarchies (Foster *et al*., 2017).

Second, we compared species richness using Simpson’s α-diversity index calculated from the phyloseq object (McMurdie & Holmes, 2013). We assessed differences of the mean indices for each heat treatment between growth stages, and their interactions using a Multi-Factor ANOVA and Tukey’s Post-Hoc test for significant groups that respected the assumptions of normality (Blakney *et al*., 2022; Blakney *et al*., 2023). Normality of the residuals was established with a Shapiro-Wilk test using the `shapiro.test` function, while the heteroscedascity of residuals was confirmed with using a Bartlett test, `bartlett.test` function. For significant ANOVAs, a post-hoc Tukey’s Honest Significant Difference test, TukeyHSD, was used to determine which groups were statistically different.

### 7. Identification of differently abundant ASV’s and specific indicator species

To refine our understanding of the abundance and composition of the *Vitis* sp. fungal communities, we used two complementary methods to identify taxa specific to growth stages and treatments. First, taxa cluster maps were used to calculate the differential abundance of ASVs between experimental groups using the `heat_tree_matrix` function from the `metadcoder` package (Foster *et al*., 2017). Second, indicator species analysis was used to detect ASVs that were preferentially abundant in pre-defined environmental groups (compartments, growth stages, treatments) using the `multiplatt` function from the `indicspecies` package (De Cáceres & Legendre, 2009), with an FDR correction. A significant indicator value is obtained if an ASV has a large mean abundance within a group, compared to another group (specificity), and has a presence in most samples of that group (fidelity) (De Cáceres & Legendre, 2009; Legendre & Legendre, 2012). The fidelity component complements the differential abundance approach between taxa clusters, which only considers abundance.

### 8. β-diversity of the *Vitis* sp. fungal communities in the phyllosphere and carposphere

To test for significant differences between the *Vitis* sp. fungal communities at different growth stages and heat treatments, we used the non-parametric permutational multivariate ANOVA (PERMANOVA), where any variation in the ordinated data distance matrix is divided among all the pairs of specified experimental factors. The PERMANOVA was calculated using the adonis `adonis` function from the `vegan` package (Oksanen *et al*., 2020), with a distance matrix calculated with the Bray-Curtis formula, 9999 permutations, and the experimental blocks were included as “strata”. We confirmed that our data met the assumption of homogeneity using the `betadispr` function, and tested using an ANOVA and Tukey’s Honest Significant Difference post-hoc test, to determine that none of the groups were statistically different.

Similarity between communities was also tested and visualized using principal co-ordinate analysis (PCoA, Legendre & Legendre, 2012) using the Bray-Curtis distance matrix. Singleton ASVs were removed before the phyloseq data were transformed using Hellinger’s transformation, such that ASVs with high abundances and few zeros are treated equivalently to those with low abundances and many zeros (Legendre & de Cáceres, 2013).

To further characterize what ecological mechanisms may be responsible for changes to the β-diversity, we partitioned β-diversity into 2 components: turnover (i.e. species replacement) and nestedness (i.e. loss/gains that result in poor species richness being a subset of richer sites; Blakney *et al*., 2024). We compared the β-diversity components for each heat treated plot at each growth stage with its cognate plot at the following growth stage (e.g. control phyllosphere at EL-32 was compared to control phyllosphere at EL-36), using the `betapart.core.abund` function from the `betapart` package (Baselga *et al*., 2023). Within the phyllosphere and the carposhere, we separately identified any significant differences among the means of each component between growth stages for each heat treatment, and their interactions, with a Multi-Factor ANOVA, as described above for α-diversity, as normality was respected.

## Results

### 1. Illumina MiSeq yielded similar numbers of fungal ASVs from both *Vitis* sp. phyllosphere and carposphere

Illumina’s MiSeq produced 1 616 465 raw reads for the whole fungal ITS dataset, which were processed through cutadapt and DADA2 (Callahan *et al*., 2016a & 2016b), where 1 160 088 reads were retained from all the experimental samples (Fig. S1). The number of ITS reads were similar between both leaf and fruit samples, and across growth stages, with a mean of 8044 ±2328 reads among leaves, and 7865 ±2281 reads among fruit (Table 1). From this, a total of 655 distinct fungal ASVs were inferred. The majority of reads from across the dataset were identified as *Dothideomycetes* (Phylum *Ascomycota*; Fig. 1 & S3). However, we also observed a consistent trend concerning the number of unique ASVs across the dataset, with a mean of 29 ±8 unique ASVs among leaf samples and a mean of 22 ±6 ASVs among fruit samples (Table 1).

**Figure 1.**
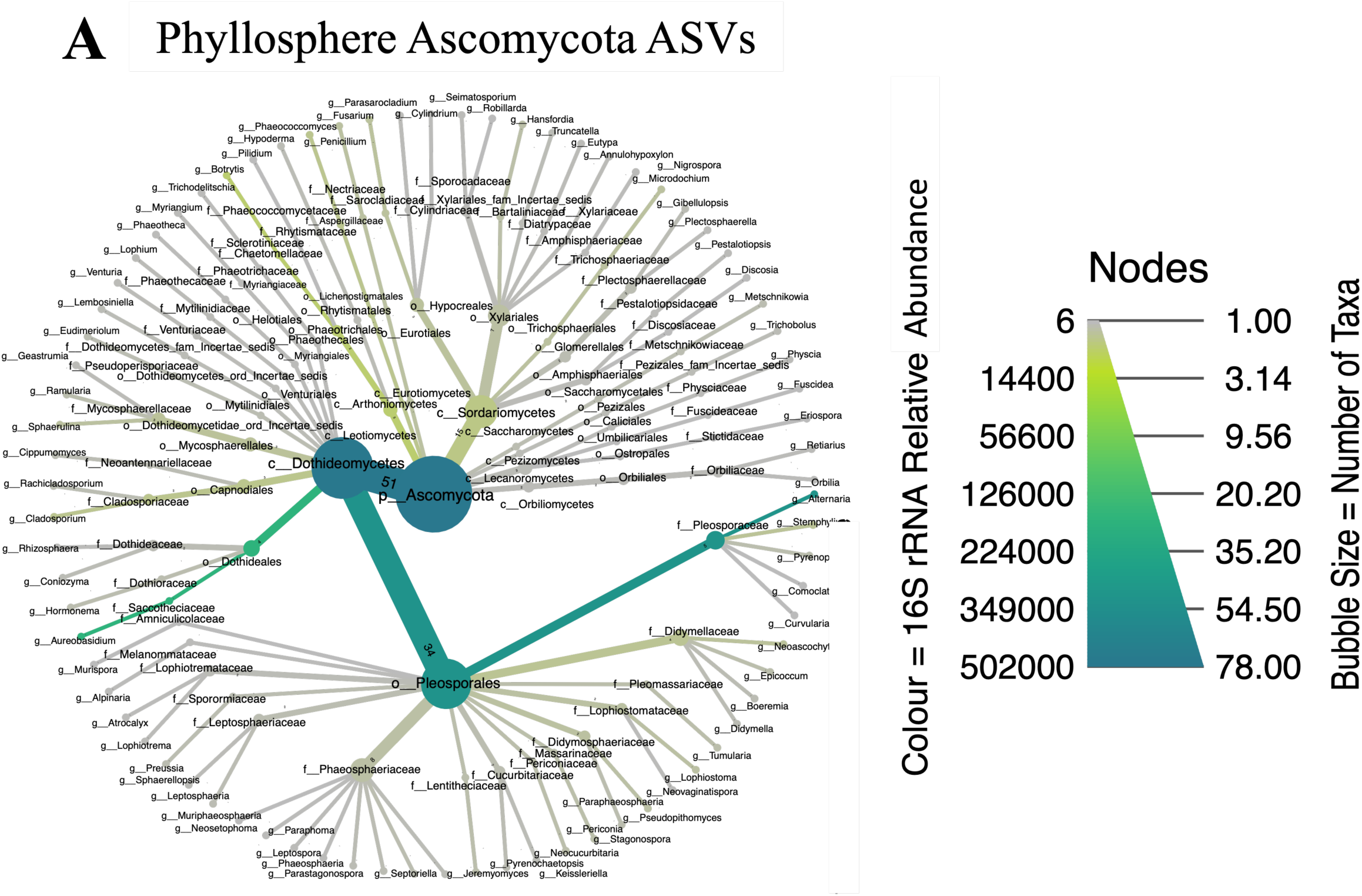
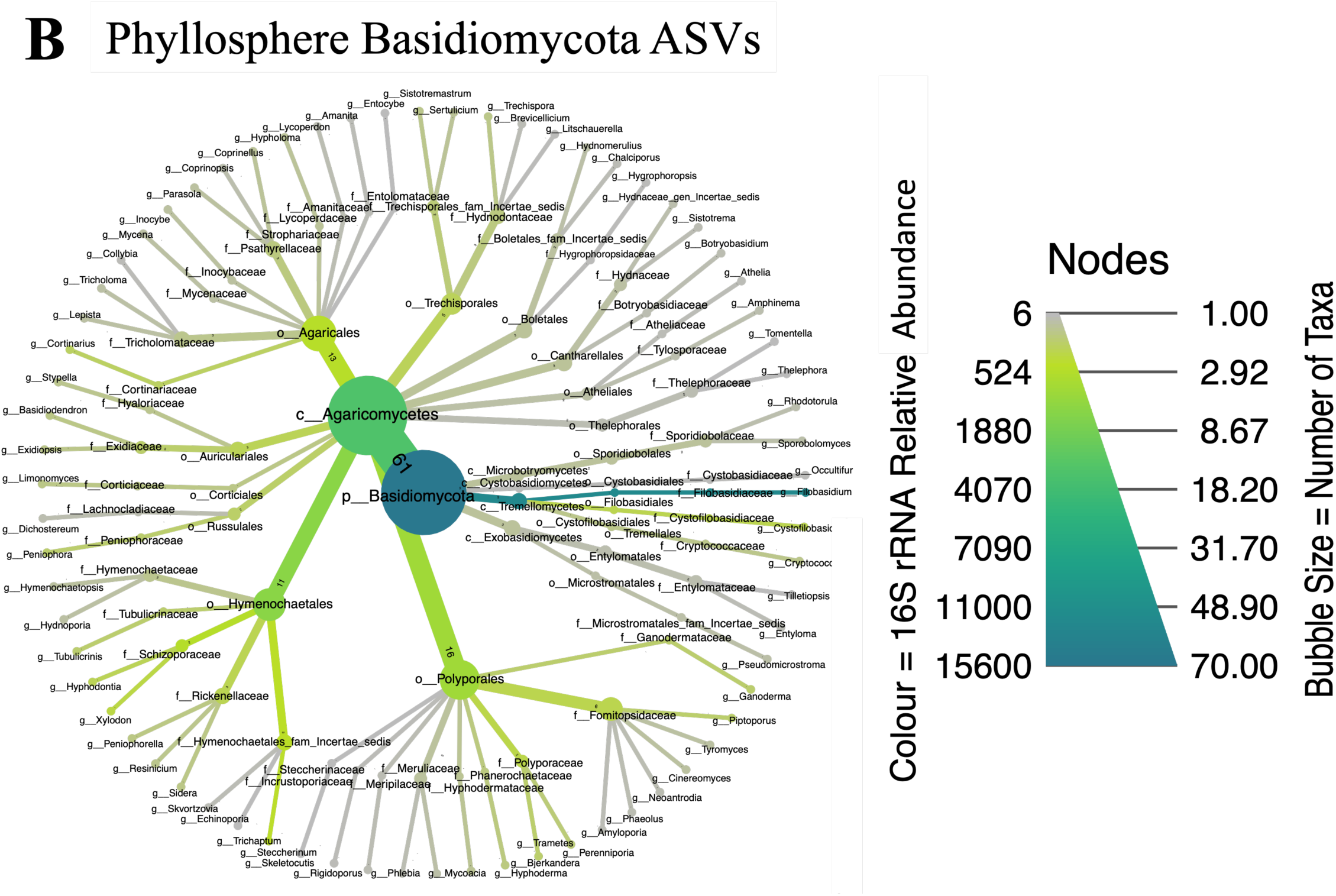
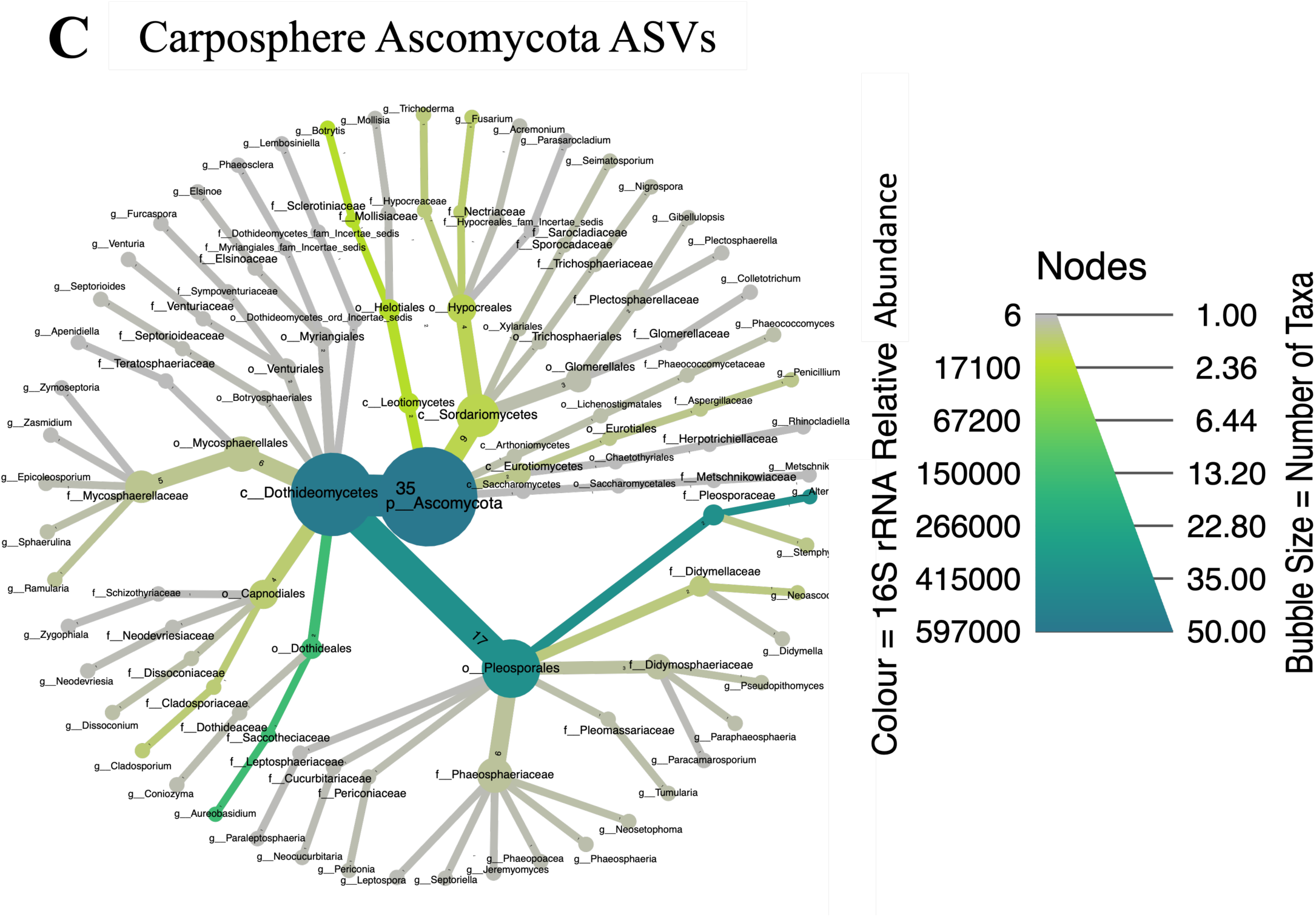
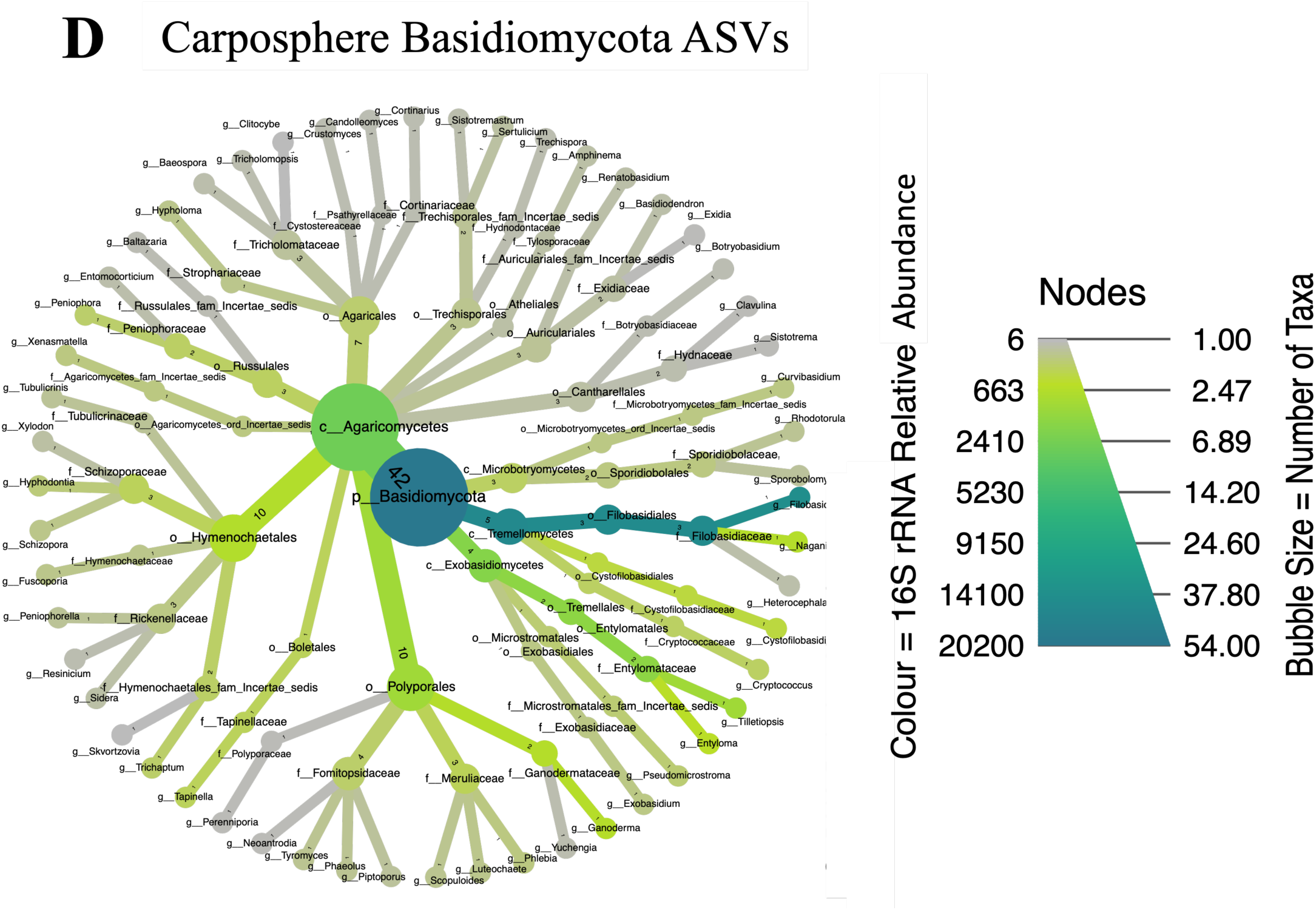
Fungal community composition from the phyllosphere (A & B) and carposphere (C & D) of *Vitis* sp. cv. L’Acadie blanc sampled across the 2020 season in Wolfville, Nova Scotia. High-quality MiSeq reads from the ITS amplicons were retained through the DADA2 pipeline, inferred as amplicon sequence variants (ASVs), and assigned taxonomy using the UNITE database. The taxa clusters illustrate that the phyllosphere communities (A & B) had higher diversity than the carposphere (C & D). Communities were dominated by Ascomycetes (A & C), of the class *Dothideomycetes*. The dominant *Dothideomycetes* were primarily *Alternaria* spp. and *Aureobasidium* spp. in both phyllosphere (A) and carposphere (C), while the dominant Basidiomycetes in the phyllosphere (B) and carposphere (D) was *Filobasidium* spp.

**Table 1.** The leaves and fruits of *Vitis* sp. cv. L’Acadie Blanc, sampled throughout the 2020 growing season in Wolfville, Nova Scotia, yielded 1 616 465 raw reads for the fungal ITS data via Illumina’s MiSeq platform at Génome Québec. The raw reads were processed through DADA2, to retain 1 160 088 reads (ITS Reads reported here) for ASV inference. A total of 655 fungal ASVs were identified across the dataset.

| Growth Stage | Compartment | ITS Reads | Fungal ASV Occurrence |
| --- | --- | --- | --- |
| EL-32 | Leaf<br>n = 13 | 6628 ±1858 | 26 ±9 |
|  | Fruit<br>n = 20 | 7225 ±1967 | 20 ±7 |
| EL-36 | Leaf<br>n = 19 | 7860 ±1901 | 28 ±5 |
|  | Fruit<br>n = 20 | 7470 ±1751 | 22 ±7 |
| EL-37 | Leaf<br>n = 18 | 9367 ±2644 | 30 ±6 |
|  | Fruit<br>n = 20 | 8980 ±2978 | 24 ±5 |
| EL-38 | Leaf<br>n = 16 | 7922 ±2137 | 33 ±9 |
|  | Fruit<br>n = 20 | 7786 ±1982 | 21 ±5 |

### 2. Fungal communities were significantly different between phyllosphere and carposphere

Overall, the PERMANOVA supported that the fungal communities from the phyllosphere and carposphere of *Vitis* sp. were significantly different (PERM *R^2^* = 0.01461, *p* = 0.044; Table 2). The interaction between growth stage and compartments (i.e. phyllosphere and carposphere) was also significant (Table 2). Several ASVs were identified as specific to the phyllosphere, though weakly significant; the ascomycetes *Nigrospora* spp. (*Trichosphaeriaceae*), *Sphaerulina* spp. (*Mycosphaerellaceae*), *Alternaria* spp. (*Pleosporaceae*), an unknown *Didymellaceae* (*Pleosporales*), and a sole basidiomycete, *Piptoporus* spp. (*Fomitopsidaceae*, Table 3). Only ASVs identified as *Cladosporium* spp. (*Cladosporiaceae*) were enriched (i.e. more reads) in the phyllosphere, compared to the carposphere (*p* < 0.05; Fig. S4). In the carposphere, only the *Polyporales* and *Hymenochetales* orders were enriched, compared to the phyllosphere (*p* < 0.05; Fig. S4), but no family or genus was found to be significant. Two ASVs were identified as specific to the fruits, the basidiomycetes *Naganishia* spp. (*Filobasidiaceae*), and *Tilletiopsis* spp. (*Entylomatales*; Table 3).

**Table 2.** PERMANOVA identified compartments, growth stage and mini-greenhouse treatments as significant experimental factors for the fungal communities harvested in 2020 from *Vitis* sp. in Wolfville, Nova Scotia. Only significant interactions are presented. PERMANOVA was calculated using a Bray-Curtis distance matrix, with 9999 permutations.

|  | ITS |  |  |
| --- | --- | --- | --- |
|  | F Model | R<sup>2</sup> | Pr (> F) |
| Compartment<sup>a</sup> | 2.4943 | 0.01461 | **0.044** |
| Growth Stage<sup>b</sup> | 1.8943 | 0.03329 | **0.027** |
| Treatment<sup>c</sup> | 3.2973 | 0.05794 | **0.001** |
| Compartment ~<br>Growth Stage | 2.0629 | 0.03625 | **0.022** |
| Compartment ~<br>Growth Stage ~<br>Treatment | 2.0613 | 0.10866 | **0.003** |

<sup>a</sup>, phyllosphere or carposphere communities
<sup>b</sup>, growth stages: EL-32 (fruit bunching), EL-36 (intermediate fruit sugar levels), EL-37 (fruit not quite ripe), and EL-38 (ripe and harvest)
<sup>c</sup>, heat treatments applied pre-veraison (before ripening), post-veraison, all-season, or untreated control

**Table 3.** Indicator fungal species were identified exclusively among the phyllosphere or carposphere of *Vitis* sp. grown during the 2020 season in Wolfville, Nova Scotia. Indicator species analysis relies on abundance and site specificity to statistically test each ASV, which we report here as a tendency (*p* < 0.1), with a FDR correction.

| | Closest Taxon | Compartment<sup>a</sup> ($p < 0.1$) |
| --- | --- | --- |
| Ascomycetes | *Alternaria* spp. | Phyllosphere |
|  | *Didymellaceae* | Phyllosphere |
|  | *Nigrospora* spp. | Phyllosphere |
|  | *Sphaerulina* spp. | Phyllosphere |
| Basidiomycetes | *Piptoporus* spp. | Phyllosphere |
|  | *Naganishia* spp. | Carposphere |
|  | *Tilletiopsis* spp. | Carposphere |

<sup>a</sup>, phyllosphere or carposphere communities

Contrary to our initial hypothesis, where we predicted that fungal diversity would increase over the growing season, we observed little influence of the different growth stages on α- (Fig. 2) and β-diversity (Fig. 3), even though we did find that different growth stages had a significant effect on the fungal communities in the phyllosphere and carposphere (PERM *R^2^* = 0.03329, *p* = 0.027; Table 2). We also found that species turnover across growth stages was more significant in explaining the β-diversity of phyllosphere and carposphere communities than species nestedness (*p.* adj < 0.001; Fig. 3D). We did not detect any taxonomic enrichments, or depletions, according to growth stage (Fig. 4), and the fungal communities remained largely taxonomically similar through time (Fig. 3A).

**Figure 2.**
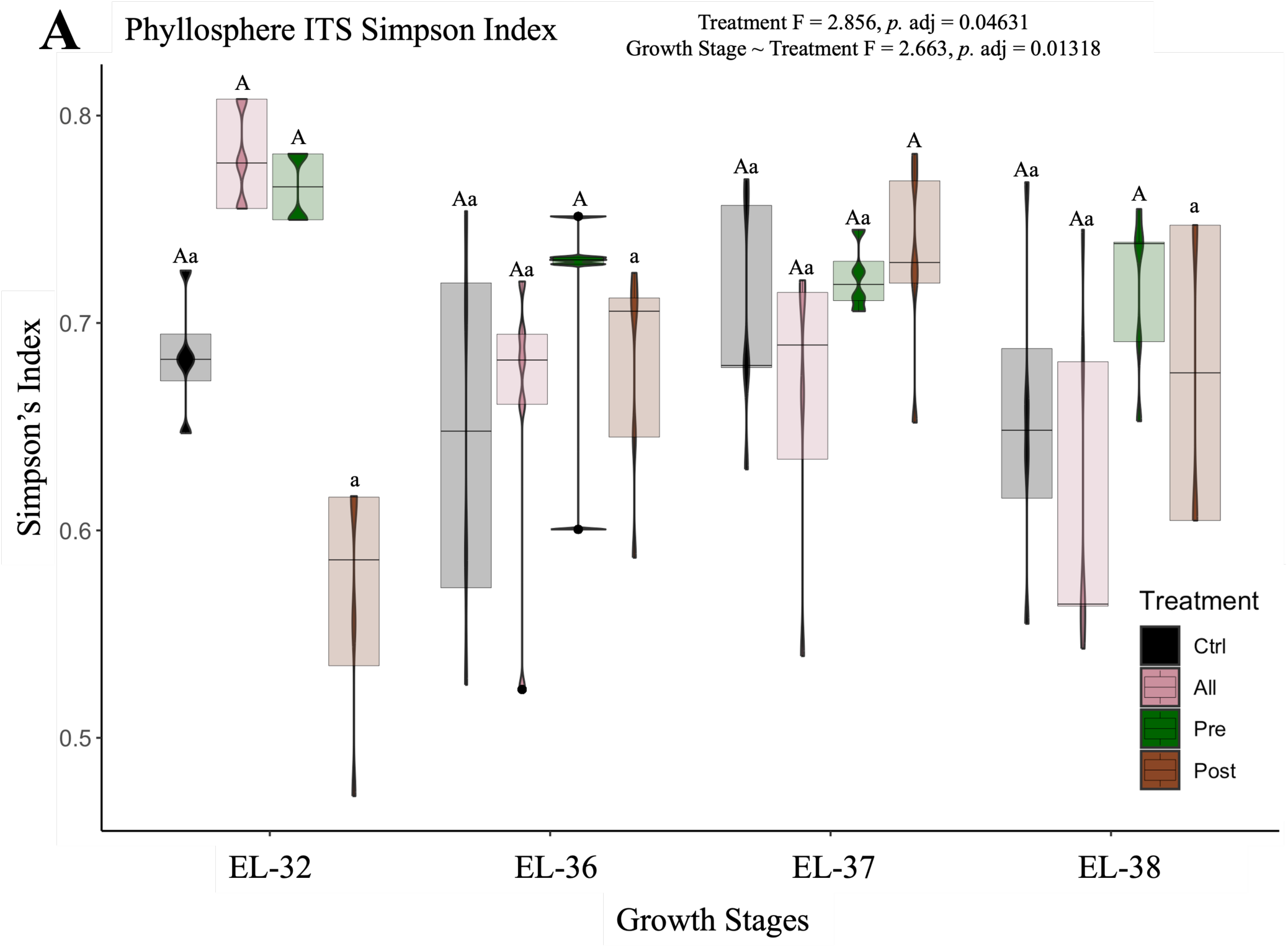
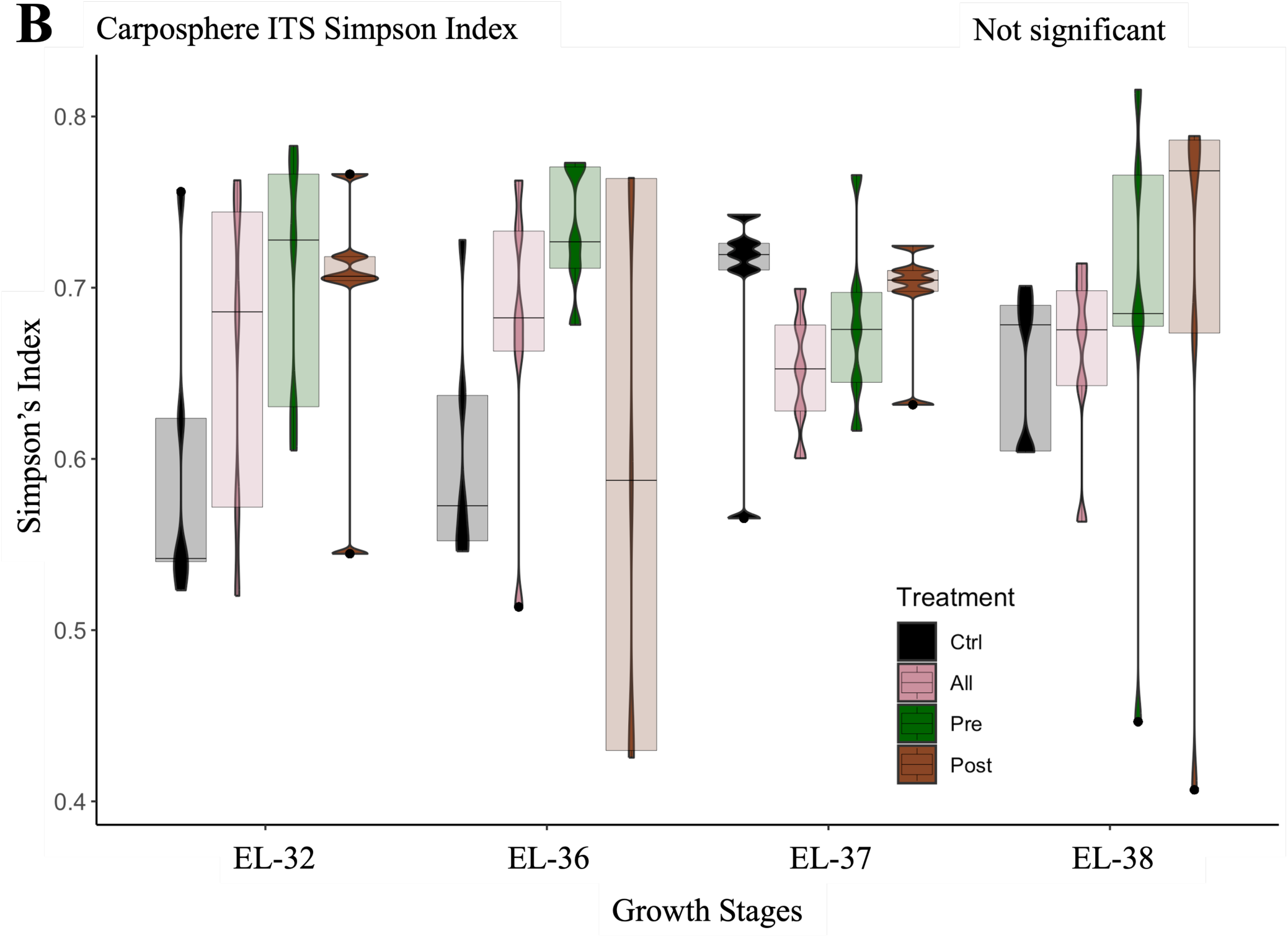
Fungal community diversity among *Vitis* sp. phyllospheres was significantly impacted by the different mini-greenhouse treatments (A, *p* = 0.04631), while the carposphere communities remained stable (B) across the 2020 season in Wolfville, Nova Scotia. (A) Simpson’s diversity index was significantly higher for phyllosphere communities treated pre-veraison across the EL-32, 36, & 38 growth stages, when compared to communities treated post-veraison. Although the growth stages were not significant factors in the model, the growth stage ∼ treatment interaction was (*p* = 0.01318). (B) Diversity was not significantly impacted by growth stage, or treatments, in the carposphere fungal communities. Diversity across growth stages and with different treatments was tested with a Multi-Factor ANOVA, where Tukey’s post-hoc test identified the statistically significant groups.

**Figure 3.**
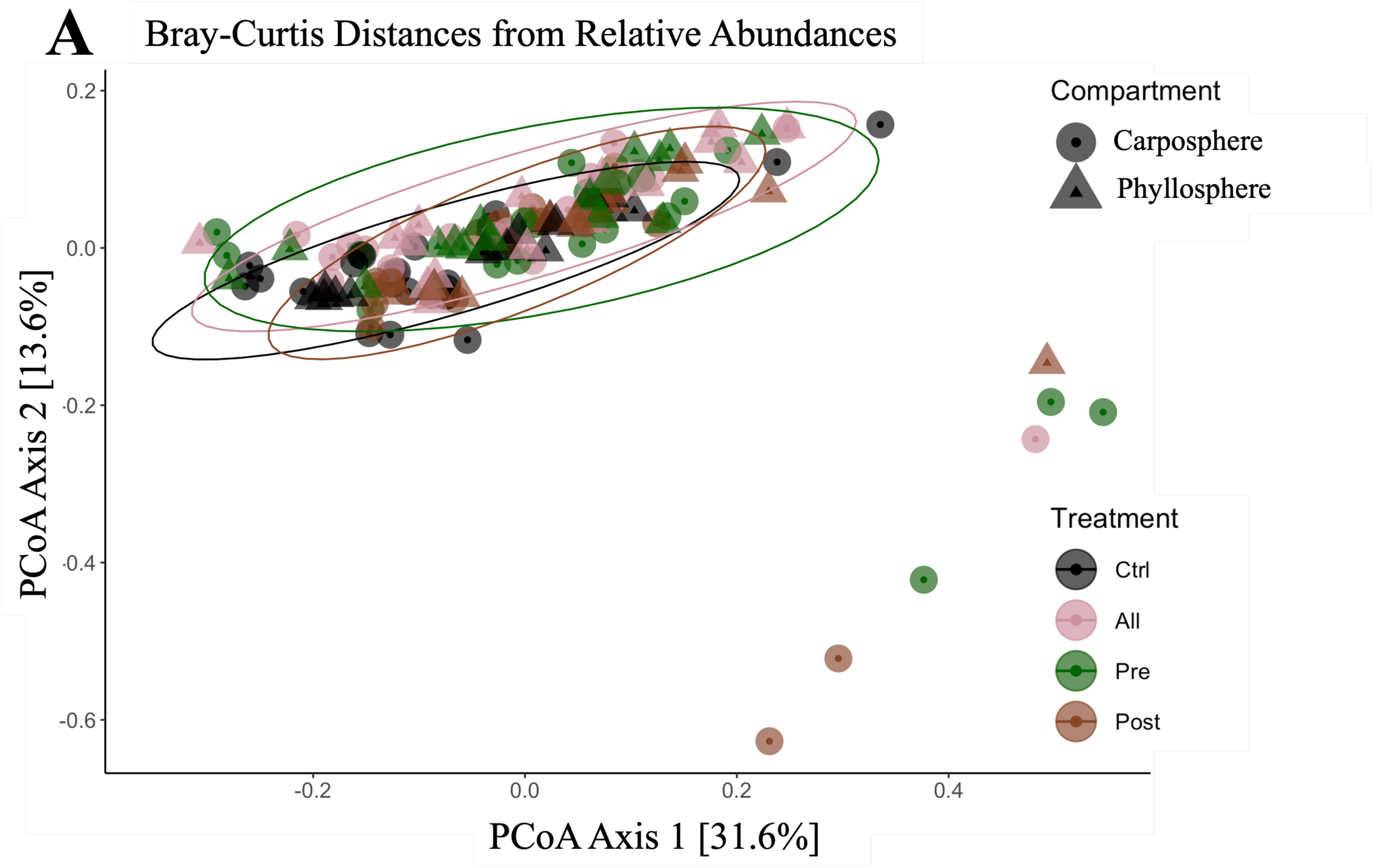
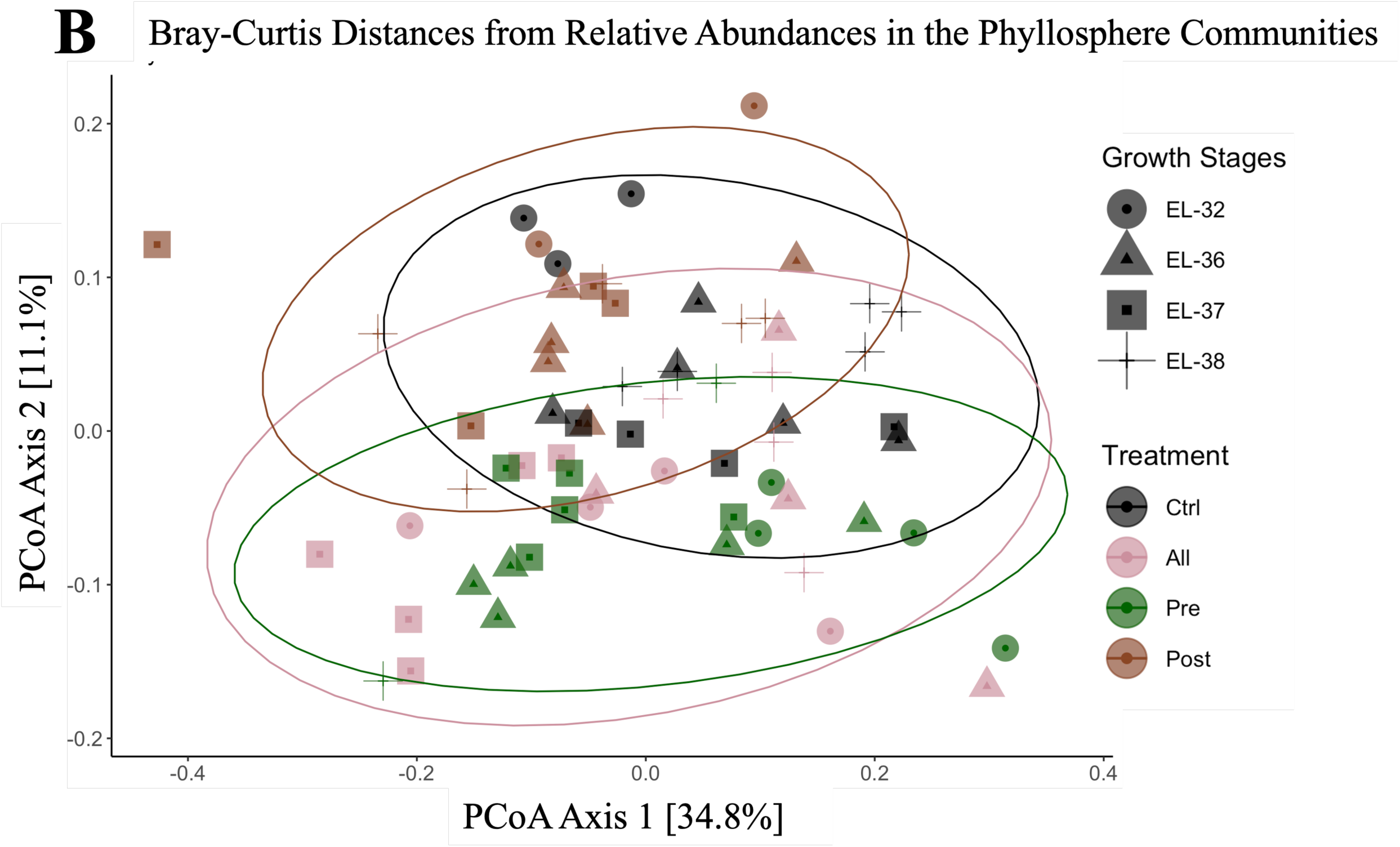
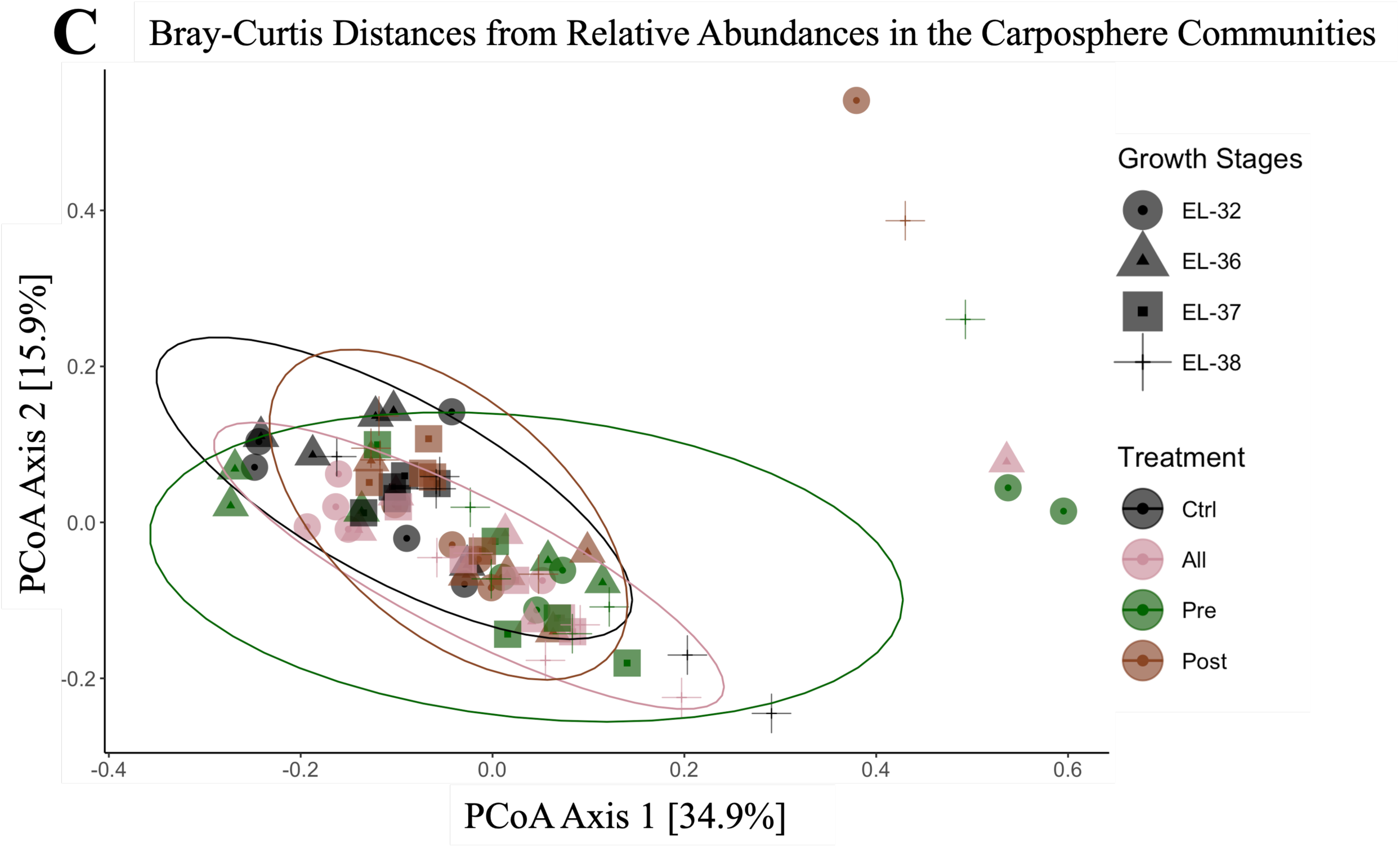
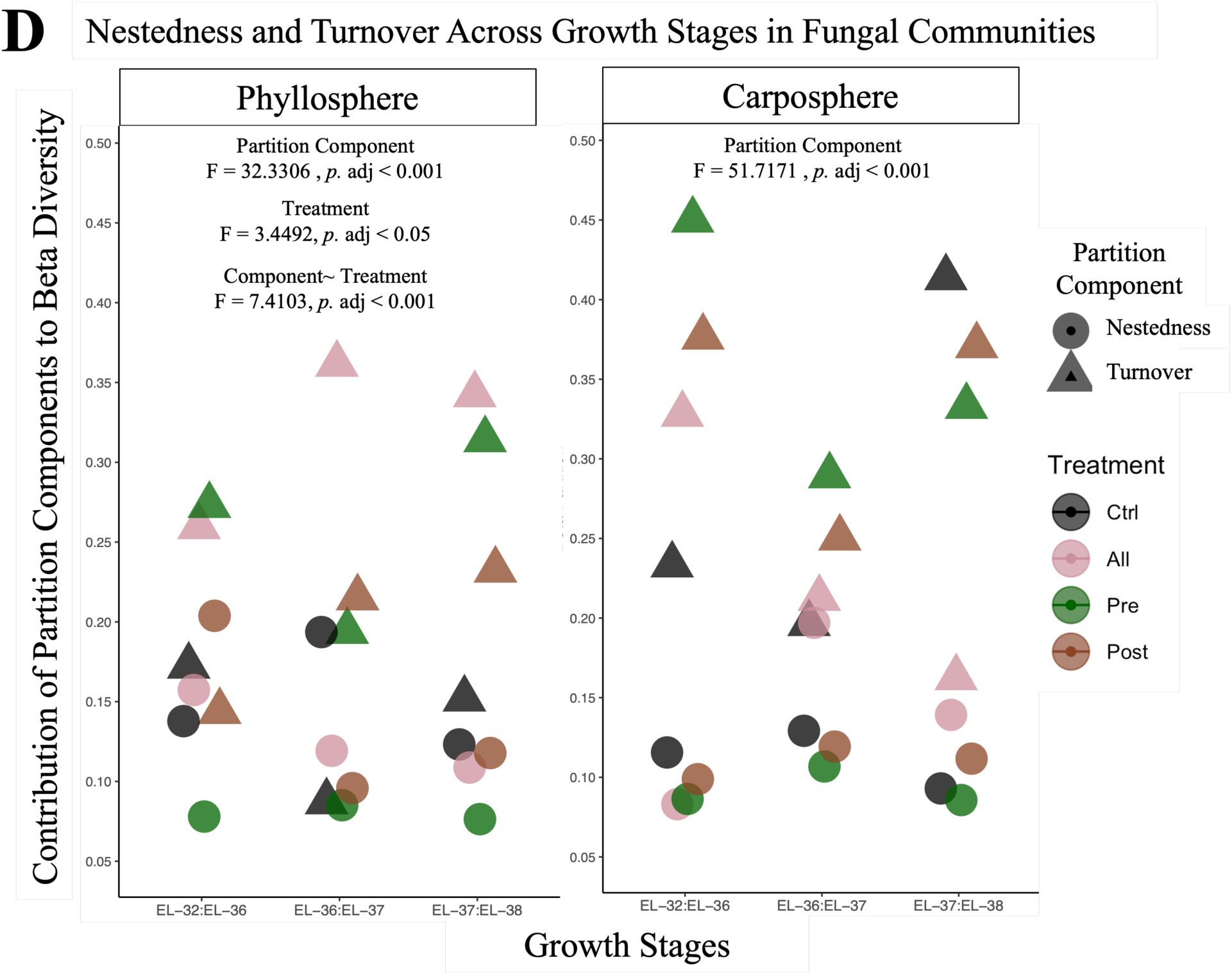
β-diversity of fungal communities identified from the phyllosphere and carposphere of *Vitis* sp. illustrates similar composition and structure between compartments and growth stages, primarily explained by species turnover. Samples were harvested throughout the 2020 growing season in Wolfville, Nova Scotia. A) Principal co-ordinate analysis captured 45.2% of the variability among the phyllosphere and carposphere communities, which remained similar in composition and diversity between compartments and across temperature treatments. (B) Principal co-ordinate analysis captured 45.9% of the variability among the phyllosphere communities, where samples from the control and post-veraison treatments were more compositionally similar compared to communities from other treatments. Phyllosphere communities from the EL-38 growth stage were also more compositionally similar, regardless of treatment, compared to communities from other growth stages. (C) Principal co-ordinate analysis captured 50.8% of the variability among the carposphere communities, which remained similar in composition and diversity across growth stages and between temperature treatments. (D) Species turnover was significantly higher than nestedness in both phyllosphere and carposphere communities (*p* < 0.001), with turnover being significantly higher among treated phyllosphere communities compared to control communities (adj. *p* < 0.05). Influence of turnover and nestedness across growth stages and with different treatments was tested with a Multi-Factor ANOVA, where Tukey’s post-hoc test identified the statistically significant groups.

**Figure 4.**
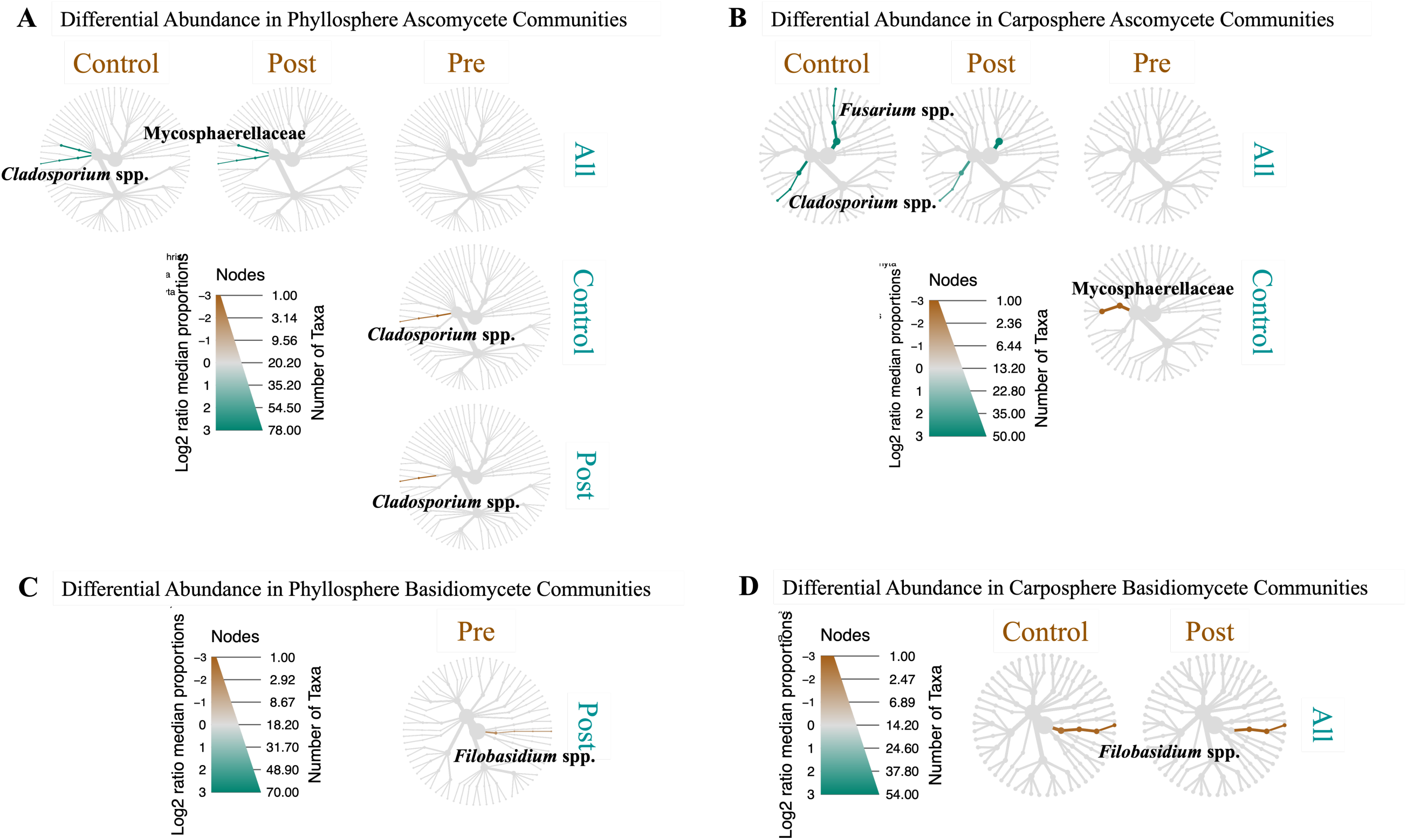
Certain fungal ASVs were significantly enriched (*p* < 0.05) in the phyllosphere (A & C) and carposphere (B & D) due to the increased temperature treatments, but not due to growth stage of *Vitis* sp. grown during the 2020 season in Wolfville, Nova Scotia. (A) Among the phyllosphere Ascomycete communities, ASVs identified as *Cladosporium* spp. and *Mycosphaerellaceae* were over-represented in samples treated all season, compared to both control and post-veraison samples. Similarly, samples treated with increased heat during pre-veraison were also enriched in *Cladosporium* spp., compared to control and post-veraison samples. (B) Among the carposphere Ascomycete communities, ASVs for *Cladosporium* and *Fusarium* spp. were enriched in samples treated all season, compared to both control and post-veraison samples. Pre-veraison carposphere samples were also enriched in the *Mycosphaerellaceae*, compared to the control samples. Among the Basidiomycete communities, phyllosphere samples treated pre-veraison were over-abundant in *Filobasidium* spp. compared to the samples treated post-veraison (C), while *Filobasidium* spp. were also enriched in the carposphere control and post-veraison samples, compared to those treated with increased heat all season (D). Significantly enriched taxa, labelled in bold, were tested between each pair of treatments. Taxa that were significantly more abundant are highlighted brown or green, following the labels for each compared host. These differential taxa clusters identified significantly enriched (i.e., abundant), using the non-parametric Kruskal test, followed by the post-hoc pairwise Wilcox test, with an FDR correction.

### 3. Increased heat significantly altered fungal communities in the phyllosphere

The different heat treatments did significantly influence the fungal communities of the phyllosphere and carposphere of *Vitis* sp (PERM *R^2^* = 0.05794, *p* < 0.001; Table 2). The interaction between treatment and growth stage was also significant (Table 2), though likely driven by the significant changes between the phyllosphere and carposphere communities, and the significant impact of the heat treatments. Although the structure and composition of the phyllosphere and carposphere communities across all treatments remained largely similar (Fig. 3A), among the fungal phyllosphere, the heat treatments had significant effects on the α-diversity. Simpson’s index was higher for phyllosphere communities treated with increased heat pre-veraison at growth stages EL-32, 36, and 38, when compared to communities treated with increased heat post-veraison (*p.* adj = 0.04631; Fig. 2A), in opposition to our initial prediction. Moreover, it is worth noting that the untreated control phyllosphere samples remained stable in terms of composition, relative abundance and diversity, throughout the experiment (Figs. 2A, 3B & 3D), contrary to our prediction. This further supports the relatively weak influence of the different growth stages on phyllosphere community structure (Fig. 2A). In the β-diversity analyses of the phyllosphere, the PCoA captured 45.9% of the variation and illustrated a shift in fungal community similarity, where communities from the control and post-veraison heated communities were more similar in composition versus communities that were heated all season and pre-veraison (Fig. 3B). The increased heat treatments also increased species turnover relative to the controls in the phyllopshere communities (*p.* adj < 0.05; Fig. 3D).

A number of fungal taxa were also significantly enriched among the phyllosphere communities due to specific treatments (*p* < 0.05; Fig. 4). ASVs belonging *Cladosporium* spp. and the *Mycosphaerellaceae* were over-represented in samples treated for the whole season with increased heat, compared to samples from the control and post-veraison heat treatment (*p* < 0.05; Fig. 4A). Samples treated with increased heat pre-veraison were also over-abundant in *Cladosporium* spp. compared to control samples, or those treated with increased heat all post-veraison (*p* < 0.05; Fig. 4A). The pre-veraison samples were also enriched in *Filobasidium* spp. compared to the samples treated with increased heat post-veraison (*p* < 0.05; Fig. 4C).

In the *Vitis* sp. carposphere communities, the α- (Fig. 2B) and β-diversities (Fig. 3) remained stable across the growing season regardless of heat treatments, contrary to our predictions. However, there were slight variations among taxa in the carposphere communities: the samples treated with increased heat all season were enriched in *Cladosporium* and *Fusarium* spp., relative to the both the control and post-veraison samples (*p* < 0.05; Fig. 4B), while the pre-veraison fruit samples were enriched in the *Mycosphaerellaceae*, compared to the controls (*p* < 0.05; Fig. 4B). Finally, the control samples and those treated with increased heat pre-veraison were enriched in *Filobasidium* spp., compared to samples treat all season (*p* < 0.05; Fig. 4D).

## Discussion

Microbial communities are influenced by temperature and are expected to be impacted globally due to increased temperature variation induced by the climate crisis (Malik *et al*., 2019; Jannson & Hofmockel, 2020; Koskella, 2020; Zhu *et al*., 2021; Perreault & Laforest-Lapointe, 2022; Tiedje *et al*., 2022). This will have significant consequences on productivity and the quality of most agricultural crops, including grapevines, as microbial communities are tightly related to the terroir characteristics of wine (Bokulich *et al*., 2014; Bokulich *et al*., 2016; Belda *et al*., 2017; Liu *et al*., 2020; Zhou *et al*., 2021; Gobbi *et al*., 2022). Therefore, changes in the fungal community of the phyllosphere, or carposphere, could have negative effects by favoring pathogen populations, as well as positive effects by favoring fungi with a role in biological control, or the regulation of pathogen populations. However, despite their important role in grapevine health and disease, the fungal community in the grapevine phyllosphere have been relatively understudied (Perazzolli *et al*., 2014; Singh *et al*., 2018; Singh *et al*., 2019; Perazzolli *et al*., 2020; Behrens & Fischer, 2022; Wicaksono *et al*., 2023; Cui *et al*., 2024), even more true for the carposphere (Kecskeméti *et al*., 2016; Deyett & Rolshausen, 2020; Testempasis *et al*., 2023). Here, our field experiment used on-the-row mini-greenhouses to increase the temperature of *Vitis* sp. cv. L’Acadie blanc across four different growth stages. We then tested how increased temperatures significantly altered the fungal communities of the phyllosphere and carposphere of *Vitis* sp. throughout the growing season. We identified the fungal communities of the phyllosphere and carposphere at each growth stage using ITS metabarcoding and measured fungal community composition and α- and β-diversity to assess the impact of the increased temperature treatments to the phyllosphere and carposphere communities.

Our results showed typical fungal communities for the phyllosphere, dominated by *Ascomycota*, with high abundance of *Dothideomycetes* (Fig. 1 & S3), primarily *Alternaria* spp. and *Aureobasidium* spp (Kraus *et al*., 2019; Molnár *et al*., 2023). Common phyllosphere fungi, including *Alternaria* and *Nigrospora* spp. were also identified as indicator species (Table 3). We observed that the increased temperature treatments had more significant effect on fungal communities than the different growth stages (Table 2, Fig. 2, 3 & 4). Although the PERMANOVA supported significant differences between the fungal communities across growth stages (Table 2), we did not observe any clear causes in terms of composition (Fig. 1, 3 & 4). This could be attributed to fungal communities being more stable through time, unlike bacterial communities (Blakney *et al*., 2022; Blakney *et al*., 2023; Blakney *et al*., 2024b). Indeed, previous studies have also reported limited temporal effects on fungal communities (Duchicela *et al*., 2013; Gschwend *et al*., 2021). Unlike bacterial communities, which grow and shift relatively rapidly, fungal communities tend to remain more stable through time and are less affected by their hosts (Hannula *et al*., 2021). This may be due to fungi having slower growth rates and being more mobile than bacteria. Thus, in our experiment it would be reasonable not to detect vastly different fungal communities across different developmental stages.

Our most significant finding was that the increased temperature during the pre-veraison stage significantly increased the α-diversity of the phyllosphere fungal communities, and that this effect was maintained throughout the growing season (Fig. 2A). This suggests that the phyllosphere fungal communities may be more prone to changes earlier in development, nearer the time microbial communities begin colonization. For instance, the increased temperature treatment could promote the establishment of primary colonizers, or pioneer species, or create other kinds of niche space, as a priority effect, which continued to structure the community throughout the season (Chase *et al*., 2010; Hiscox *et al*., 2015; Kraus *et al*., 2019). Interestingly, increasing the temperature all-season did not yield a similar increase in α-diversity of the phyllosphere communities as the pre-veraison treatment did (Fig. 2A). Both treatments shared a similar temperature increase at a similar developmental stage for the leaf, but we did not observe a consistent impact on the diversity of the fungal communities. This could suggest that any priority effect, or additional niche space, created in the phyllosphere by the pre-veraison temperature treatment may not be sustained by the prolonged all-season heat treatment. Instead, the longer all-season heat treatment may provide more time for the fungal communities in the phyllosphere to recalibrate.

The potential pre-veraison priority effect could also be related to the significant enrichment of ASVs belonging to the *Cladosporium*, or *Filobasidium* spp. in the phyllosphere communities (Fig. 4). Both groups have the potential to be antagonistic toward the plant: *Filobasidium* spp. are common basidiomycetes and tend to be poor saprotrophs, favouring environments where they can parasitize other fungi, or plants (Weiss *et al*., 2014; Detheridge *et al*., 2020), while *Cladosporium* spp. are also well-known in the phyllosphere and diverse plant pathogens (Krause *et al*., 2019; Cosseboom & Hu, 2023). We also identified a number of ASVs specific to the phyllosphere as indicator species from among the *Dothideomycetes* class, which dominated our communities by relative abundance (Fig. 1), including *Sphaerulina* and *Alternaria* spp. (Table 3). A number of *Sphaerulina* species are known as causative agents of leaf spot disease among other plants (Ali *et al*., 2021). Equally diverse among grapevine phyllosphere communities are *Alternaria* spp., which range from saprotrophs to pathogens (Molnár *et al*., 2023). *Alternaria* spp. produce a wide range of metabolites and depending on their host and environmental factors can contribute to anti-microbial activities or virulence (Molnár *et al*., 2023).

The most common diseases present in Northern climates are downy mildew (*Plasmopara viticola*), powdery mildew (*Erisyphe necator*), anthracnose (*Elsinoe ampelina*), and Botrytis bunch rot (*Botrytis cinerea*), but there are more than 50 diseases listed in the American Compendium of Grape Disease (Carisse *et al*., 2006; Wilcox *et al*., 2015). Among the common phytopathogens identified in our data *Botrytis*, *Cladosporium*, and *Fusarium* spp., were all detected in the phyllosphere, and in higher abundance in the carposphere (Fig. 1; Lorenzini & Zapparoli, 2015; Cosseboom & Hu, 2023; Bustamante *et al*., 2024). These diseases not only affect yields, but also the quality of the grapes and, therefore, the wine they produced (Lorenzini & Zapparoli, 2015). Currently, these diseases can be controlled by the application of synthetic fungicides, which are still a key input in viticulture because most traditional wine grape varieties (*V*. *vinifera* varieties) are highly sensitive to them (Pedneault & Provost, 2016; Provost & Pedneault, 2016). However, more and more winegrowers are seeking to reduce the use of these product in favor of biological disease control along with the introduction of resistant grape varieties (Pedneault & Provost, 2016; Provost & Pedneault, 2016; Zhang *et al*., 2020; Candel *et al*., 2023; Oliver *et al*., 2024).

Biological disease control can be achieved by applying registered biological control agents, but also by promoting populations of biological control agents naturally present in the vineyard; an approach widely used in agroecology (Krause *et al*., 2019; IPPC, 2024). Our data suggests that primary colonizers could be used in biological control, or even in agroecological protection, specifically by applying a cocktail of organisms early in the season composed of good colonizers, or pioneer species, in order to limit the possible colonization by phytopathogenic organisms (Krause *et al*., 2019). Microbes—essentially bacteria, fungi and nematodes (Lindow & Brandl, 2003; Perazzolli *et al*., 2014; Sapkota *et al*., 2015; Laforest-Lapointe *et al*., 2017; Koskella, 2020)—that colonize the phyllosphere or carposphere (Vorholt JA. 2012) can act as biocontrol agents insofar as they compete for space and nutrients. For example, *Aerobasidium pullulans* and *Trichoderma* spp. compete with the grapevine pathogen *Botrytis cinerea* (Fedele *et al*., 2020); ASVs for both *Aerobasidium* and *Trichoderma* spp. were detected in our data (Fig. 1). The advantage of using organisms naturally occurring in the vineyard is their redundancy, where several can act in concert, or individually if weather conditions favor one species over another. Fungal communities can therefore play an important role in the resilience of agrosystems, including viticulture, to climate change, and as a tool for agroecological control of grapevine diseases. Our study provides a robust baseline for the succession of fungal communities across the development of the phyllosphere and carposphere that future experiments can use, particularly in determining the ecological roles of the fungal ASVs we report.

## Conclusion

Temperature variations induced by the climate crisis are a significant threat for global agricultural, including viticulture. Canadian viticulture reports being least prepared to adapt to future for temperature variation (Jobin Poirier et al., 2020). Moreover, climate changes also have important consequences for the resident above-ground fungal communities. To date, there have been very few experiments exploring the relationship between grapevines and their microbial communities, and to our knowledge, none on the hybrid grapevine cultivars grown in Atlantic Canada. We used on-the-row mini-greenhouses to test whether increases in temperature at different stages of fruit development (pre-veraison and post-veraison) or a global rise in temperature (whole season) altered the fungal communities of grapevine (*Vitis* sp. cv. L’Acadie blanc) phyllosphere and carposphere. We hypothesized that increased heat treatments would significantly increase fungal diversity of the phyllosphere and carposphere regardless of development stage. As such, leaves and fruits were treated with increased heat, sampled at four developmental stages, and the fungal communities were identified via ITS metabarcoding. We found specific phyllosphere and carposphere communities, though they tended to remain stable across development. However, we did observe significant changes in the phyllosphere communities pre-veraison due to increased temperatures. This could demonstrate a potential sustainable management solution, where on-the-row mini-greenhouses, or tunnels, could be deployed early in the growing season to foster more diverse fungal communities. This diversity may contribute to communities that are more resilient to phytopathogens. This could allow producers to avoid having to resort to more costly options, such as additional fungicides, or external heaters. This knowledge may also be integrated into decision making processes for selecting sites for new vineyards, where historic meteorological data could be used to identify sites with warmer conditions prevalent early in the season. Finally, our study provides a robust baseline for future work to determine the precise ecological roles of the fungal ASVs that were enriched due to the increased temperature, and more broadly for the adaptation of Atlantic Canadian grapevines and their fungal communities to future climates.

## Acknowledgements

AJCB performed the DNA extraction and sequencing prep, analyzed the data, and drafted the manuscript with input from all co-authors. This work was supported by a Mitacs Acceleration Grant to AJCB & KP (Grant Number IT31632), which we gratefully acknowledge. We thank graduate students Francisco Campos Arguedas and Guillaume Sarrailhé, that performed the greenhouse experiment as part of their projects, and Paméla Nicolle at Université du Québec en Outaouais for technical help.

## References

Abarenkov, K., Zirk, A., Piirmann, T., Pöhönen, R., Ivanov, F., Nilsson, R. H., & Kõljalg, U. (2020). UNITE general FASTA release for eukaryotes. Version 04.02.2020. UNITE Community.

Abarenkov, K., Zirk, A., Piirmann, T., Pöhönen, R., Ivanov, F., Nilsson, R. H., & Kõljalg, U. (2022). UNITE general FASTA release for Fungi. Version 16.10.2022. UNITE Community.

Ausseil, A. G. E., Law, R. M., Parker, A. K., Teixeira, E. I., & Sood, A. (2021). Projected wine grape cultivar shifts due to climate change in New Zealand. Frontiers in Plant Science 12:618039.

Bell, T. H., Stefani, F. O. P., Abram, K., Champagne, J., Yergeau, É., Hijri, M., & St-Arnaud, M. (2016). A diverse soil microbiome degrades more crude oil than specialized bacterial assemblages obtained in culture. Applied and Environmental Microbiology 82:5530–5541.

Baselga, A., Orme, D., Villeger, S., De Bortoli, J., Leprieur, F., Logez, M., Martinez-Santalla, S., Martin-Devasa, R., Gomez-Rodriguez, C., Crujeiras, R. (2023). betapart: partitioning beta diversity into turnover and nestedness components. R package version 1.6.

Behrens, F. H., & Fischer, M. (2022) Evaluation of different phyllosphere sample types for parallel metabarcoding of Fungi and Oomycetes in *Vitis vinifera*. Phytobiomes Journal 6:207–213.

Belda, I., Zarraonaindia, I., Perisin, M., Palacios, A., & Acedo, A. (2017) From vineyard soil to wine fermentation: microbiome approximations to explain the “terroir” concept. Frontiers in Microbiology 8:821.

Blakney, A. J. C., Bainard, L. D., St-Arnaud, M., & Hijri, M. (2022). *Brassicaceae* host plants mask the feedback from the previous year’s soil history on bacterial communities, except when they experience drought. Environmental Microbiology 24:3529–3548.

Blakney, A. J. C., Bainard, L. D., St-Arnaud, M., & Hijri, M. (2023). Soil chemistry and soil history significantly structure Oomycete communities in *Brassicaceae* crop rotations. Applied and Environmental Microbiology 89:e01314–22.

Blakney, A. J. C., Morvan, S., Lucotte, M., Moingt, M., Charbonneau, A., Bipfubusa, M., Gonzalez, E., & Pitre, F. E. (2024a) Site properties, environmental factors, and crop identify influence soil bacterial communities more than municipal biosolid application. Science of the Total Environment 926:171854.

Blakney, A. J. C., St-Arnaud, M., & Hijri, M. (2024b). Does soil history decline in influencing the structure of bacterial communities of *Brassica napus* host plants across different growth stages? ISME Communications 4:ycae019.

Bokulich, N. A., Thorngate, J. H., Richardson, P. M., & Mills, D. A. (2014) Microbial biogeography of wine grapes is conditioned by cultivar, vintage, and climate. Proceedings of the National Academy of Sciences, USA 111:E139–E148.

Bokulich, N. A., Collins, T. S., Masarweh, C., Allen, G., Heymann, H., Ebeler, S. E., & Mills D. A. (2016) Associations among wine grape microbiome, metabolome, and fermentation behavior suggest microbial contribution to regional wine characteristics. mBio 7:e00631–16.

Bonada, M, Jeffery, D. W., Petrie, P. R., Moran, M. A., & Sadras V. O. (2015) Impact of elevated temperature and water deficit on the chemical and sensory profiles of Barossa Shiraz grapes and wines. Australian Journal of Grape & Wine Research 21:240–253.

Bustamante, M. I., Todd, C., Elfar, K., Hamid, M. I., Garcia, J. F., Cantu, D., Rolshausen, P. E., & Eskalen A. (2024) Identification and pathogenicity of *Fusarium* species associated with young vine decline in California. Plant Disease 108:1053–1061.

Callahan, B. J., McMurdie, P. J., Rosen, M. J., Han, A. W., Johnson, A. J. A., & Holmes, S. P. (2016a). DADA2: High-resolution sample inference from Illumina amplicon data. Nature Methods 13:581–583.

Callahan, B. J., Sankaran, K., Fukuyama, J. A., McMurdie, P. J., & Holmes, S. P. (2016b). Bioconductor workflow for microbiome data analysis: from raw reads to community analyses. F1000Research 5:1492.

Campos-Arguedas, F., Sarrailhé, G., Nicolle, P., Dorais, M., Brereton, N. J. B., Pitre, F. E., & Pedneault, K. (2022). Different temperature and UV patterns modulate berry maturation and volatile compounds accumulation in *Vitis* sp. Frontiers in Plant Science 13:862259.

Candel, J., Pe’er, G., & Finger, R. (2023) Science calls for ambitious European pesticide policies. Nature Food 4:272.

Carisse, O., Bacon, R., Lasnier, J., & McFadden-Smith, W. (2006) Identification Guide to the Major Diseases of Grapes. Editor Agriculture and Agri-Food Canada, Publication 10092E.

Chase, J. M. (2010) Stochastic community assembly causes higher biodiversity in more productive environments. Science 328:1388–1391.

Coombe, B. G. (1995). Growth stages of the grapevine: adoption of a system for identifying grapevine growth stages. Australian Journal of Grape & Wine Research 1:104–110.

Cosseboom, S. D., & Hu, M. (2023) Identification and pathogenicity of *Cladosporium*, Fusarium, and Diaporthe spp. associated with late-season bunch rots of grape. Plant Disease 107:2929–2934.

Cui, S., Zhou, L., Fang, Q., Xiao, H., Jin, D., & Liu, Y. (2024) Growth period and variety together drive the succession of phyllosphere microbial communities of grapevine. Science of The Total Environment 950:175334.

De Cáceres, M., & Legendre, P. (2009). Associations between species and groups of sites: indices and statistical inference. Ecology 90(12):3566–3574.

Delavaux, C. S., Bever, J. D., Karppinen, E. M., & Bainard, L. D. (2020). Keeping it cool: soil sample cold pack storage and DNA shipment up to 1 month does not impact metabarcoding results. Ecology Evolution 00:1–13.

Detheridge, A. P., Cherrett, S., Clasen, L. A., Medcalf, K., Pike, S., Griffith, G. W., & Scullion, J. (2020). Depauperate soil fungal populations from the St. Helena endemic *Commidendrum robustum* are dominated by *Capnodiales*. Fungal Ecology 45:100911.

Deyett, E., & Rolshausen, P. E. (2020) Endophytic microbial assemblage in grapevine. FEMS Microbiology Ecology 96:fiaa053.

Di Paola, M., Gori, A., Stefanini, I., Meriggi, N., Renzi, S., Nenciarini, S., Cerasuolo, B., Moriondo, M., Romoli, R., Pieraccini, G., Baracchi, D., Turillazzi, F., Turillazzi, S., & Cavalieri, D. (2023). Using wasps as a tool to restore a functioning vine grape mycobiota and preserve the mycobial “terroir”. Scientific Reports 13:16544.

Duchicela, J., Sullivan, T. S., Bontti, E., & Bever, J. D. (2013). Soil aggregate stability increase is strongly related to fungal community succession along an abandoned agricultural field chronosequence in the Bolivian Altiplano. Journal of Applied Ecology 50:1266–1273.

Fedele, G., Brischetto, C., & Rossi, Vi. (2020) Biocontrol of *Botrytis cinerea* on grape berries as influenced by temperature and humidity. Frontiers of Plant Science 11:1232.

Foster, Z., Sharpton, T., & Grünwald, N. (2017). Metacoder: An R package for visualization and manipulation of community taxonomic diversity data. PLoS Computational Biology 13:1–15.

Gobbi, A., Acedo, A., Imam, N., Santini, R. G., Ortiz-Álvarez, R., Ellegaard-Jensen, L., Belda, I., & Hansen, L. H. (2022) A global microbiome survey of vineyard soils highlights the microbial dimension of viticultural *terroirs*. Communications Biology 5:241.

Gschwend, F., Hartmann, M., Hug, A. S., Enkerli, J., Gubler, A., Frey, B., Meuli, R. G., & Widmer, F. (2021). Long-term stability of soil bacterial and fungal community structures revealed in their abundant and rare fractions. Molecular Ecology 30:4305–4320.

Hannula, S.E., Heinen, R., Huberty, M., Steinauer, K., De Long, J.R., Jongen, R., et al. (2021) Persistence of plant-mediated microbial soil legacy effects in soil and inside roots. Nature Comms 12:5686.

He, L., Mazza Rodrigues, J. L., Soudzilovskaia, N. A., Barceló, M., Olsson, P. A., Song, C., Tedersoo, L., Yuan, F., Yuan, F., Lipson, D. A., & Xu, X. (2020). Global biogeography of fungal and bacterial biomass carbon in topsoil. Soil Biology and Biochemistry 151:108024.

Hiscox, J., Savoury, M., Müller, C., Lindahl, B. D., Rogers, H. J., & Brody, L. (2015). Priority effects during fungal community establishment in beech wood. ISME Journal 9:2246– 2260.

IPPC (2024) Glossary of phytosanitary terms. International standard for phytosanitary measures (ISPM) Number 5. International Plant Protection Committee. Food and Agriculture Organization (FAO), Bangkok, Rome.

Jansson, J.K., & Hofmockel, K.S. (2020) Soil microbiomes and climate change. Nature Reviews Microbiology 18:35–46.

Kecskeméti, E., Berkelmann-Löhnertz, B., & Reineke, A. (2016) Are epiphytic microbial communities in the carposphere of ripening grape clusters (*Vitis vinifera* L.) different between conventional, organic, and biodynamic Grapes? PLoS One 11:e0160852.

Koskella, B. (2020) The phyllosphere. Current Biology 30:R1096–R1169.

Kraus, C., Voegele, R. T., & Fischer, M. (2019) Temporal development of the culturable, endophytic fungal community in healthy grapevine branches and occurrence of GTD-associated fungi. Microbial Ecology 77:866–876.

Laforest-Lapointe, I., Paquette, A., Messier, C., & Kembel, S. W. (2017). Leaf bacterial diversity mediates plant diversity and ecosystem function relationships. Nature 546:145–147,

Lay, C. Y., Bell, T. H., Hamel, C., Harker, K. N., Mohr, R., Greer, C. W., Yergeau, É., & St-Arnaud, M. (2018). Canola root–associated microbiomes in the Canadian prairies. Frontiers in Microbiology 9:1–19.

Leff, J. W., Bardgett, R. D., Wilkinson, A., Jackson, B. G., Pritchard, W. J., De Long, J. R., Oakley, S., Mason, K. E., Ostle, N. J., Johnson, D., Baggs, E. M., & Fierer, N. (2018). Predicting the structure of soil communities from plant community taxonomy, phylogeny, and traits. ISME Journal 12:1794–1805.

Legendre, P., & De Cáceres, M. (2013). Beta diversity as the variance of community data: dissimilarity coefficients and partitioning. Ecology Letters 16:951–963.

Legendre, P., & Legendre, L. (2012). Numerical Ecology. 3rd ed. Elsevier. Amsterdam, The Netherlands.

Lesk, C., Anderson, W., Rigden, A., Coast, O., Jägermeyr, J., McDermid, S., Davis, K. F., & Konar, M. (2022). Compound heat and moisture extreme impacts on global crop yields under climate change. Nature Reviews Earth & Environment 3:872–889.

Lindow, S.E., & Brandl, M. T. (2003) Microbiology of the phyllosphere. Applied and Environmental Microbiology 69:1875–1883.

Liu, D., Chen, Q., Zhang, P., Chen, D., & Howell, K. S. (2020) The fungal microbiome is an important component of vineyard ecosystems and correlates with regional distinctiveness of wine. mSphere 5:e00534-20.

Lorenzini, M., & Zapparoli, G. (2015) Occurrence and infection of *Cladosporium*, *Fusarium*, *Epicoccum* and *Aureobasidium* in withered rotten grapes during post-harvest dehydration. Antonie Van Leeuwenhoek 108:1171–80

Malik, A. A., Martiny, J. B. H., Brodie, E. L., Martiny, A. C., Treseder, K. K., & Allison, S. D. (2020) Defining trait-based microbial strategies with consequences for soil carbon cycling under climate change. ISME Journal 14:1–9.

Martin, M. (2011). Cutadapt removes adapter sequences from high-throughput sequencing reads. EMBnet.Journal 17:10–12.

Masson-Delmotte, V., Zhai, P., Pirani, A., Connors, S. L., Péan, C., Berger, S., et al. (2021). Climate Change 2021: The Physical Science Basis. Contribution of Working Group I to the Sixth Assessment Report of the Intergovernmental Panel on Climate Change. IPCC, Geneva.

McMurdie P, & Holmes S (2013) Phyloseq: an R package for reproducible interactive analysis and graphics of microbiome census data. PLoS One 8(4):e61217.

Molnár, A., Knapp, D. G., Lovas, M., Tóth, G., Boldizsár, I., Váczy, K. Z., & Kovács, G. M. (2023) Untargeted metabolomic analyses support the main phylogenetic groups of the common plant-associated *Alternaria* fungi isolated from grapevine (*Vitis vinifera)*. Scientific Reports 13:19298.

Morvan, S., Meglouli, H., Lounès-Hadj Sahra, A., & Hijri, M. (2020). Into the wild blueberry (*Vaccinium angustifolium*) rhizosphere microbiota. Environmental Microbiology 22: 3803–3822.

Nottingham, A. T., Fierer, N., Turner, B. L., Whitaker, J., Ostle, N. J., McNamara, N. P., Bardgett, R. D., Leff, J. W., Salinas, N., Silman, M. R., Kruuk, L. E. B., & Meir, P. (2018). Microbes follow Humboldt: temperature drives plant and soil microbial diversity patterns from the Amazon to the Andes. Ecology 99:2455–2466.

Oksanen, J., Blanchet, F. G., Friendly, M., Kindt, R., Legendre, P., McGlinn, D., Minchin, P. R., O’Hara, R. B., Simpson, G. L., Solymos, P., Stevens, M. H. H., Szoecs, E., & Wagner, H. (2020). Vegan: Community Ecology Package. R package version 2.5–7

Oliver, C., Cooper, M., Ivey, M. L., Brannen, B., Miles, T., Lowder, S., Mahaffee, W., & Moyer, M. M. (2024) Fungicide use patterns in select United States wine grape production regions. Plant Disease 108:104–112.

Pedneault, K., Dorais, M., & Angers, P. (2013) Flavor of cold-hardy grapes: impact of berry maturity and environmental conditions. Journal of Agricultural and Food Chemistry 61: 10418–10438

Pedneault, K., & Provost, C. (2016) Fungus resistant grape varieties as a suitable alternative for organic wine production: Benefits, limits, and challenges. Scientia Horticulturae 208:57–77.

Perazzolli, M., Antonielli, L., Storari, M., Puopolo, G., Pancher, M., Giovannini, O., Pindo, M., & Pertot, I. (2014) Resilience of the natural phyllosphere microbiota of the grapevine to chemical and biological pesticides. Applied and Environmental Microbiology 80:3585– 3596.

Perazzolli, M., Nesler, A., Giovannini, O., Antonielli, L., Puopolo, G., & Pertot, I. (2020) Ecological impact of a rare sugar on grapevine phyllosphere microbial communities. Microbiological Research 232: 126387.

Perreault, R., & Laforest-Lapointe, I. (2022) Plant-microbe interactions in the phyllosphere: facing challenges of the anthropocene. ISME Journal 16:339–345.

Jobin Poirier, E., Plummer, R., & Pickering, G. (2021) Climate change adaptation in the Canadian wine industry: strategies and drivers. The Canadian Geographer / Le Géographe Canadien 65: 368–381.

Provost, C., & Pedneault, K. (2016) The organic vineyard as a balanced ecosystem: Improved organic grape management and impacts on wine quality. Scientia Horticulturae 208:43–56.

R Core Team (2020). R: A language and environment for statistical computing. R Foundation for Statistical Computing, Vienna, Austria.

Romero-Olivares, A. L., Allison, S. D., & Treseder, K. K. (2017). Soil microbes and their response to experimental warming over time: a meta-analysis of field studies. Soil Biology and Biochemistry 107:32–40.

Roy, S., Coldren, C., Karunamurthy, A., Kip, N. S., Klee, E. W., Lincoln, S. E., Leon, A., Pullambhatla, M., Temple-Smolkin, R. L., Voelkerding, K. V., Wang, C., & Carter, A. B. (2018). Standards and guidelines for validating next-generation sequencing bioinformatics pipelines. Journal of Molecular Diagnostics 20:4–27.

Singh, P., Gobbi, A., Santoni, S., Hansen, L. H., This, P., & Péros, J.P. Assessing the impact of plant genetic diversity in shaping the microbial community structure of *Vitis vinifera* phyllosphere in the Mediterranean. Frontiers in Life Science 11:35–46.

Singh, P., Santoni, S., Weber, A., This, P., & Péros, J.P. (2019) Understanding the phyllosphere microbiome assemblage in grape species (*Vitaceae*) with amplicon sequence data structures. Scientific Reports 9:14294.

Singh, B. K., Delgado-Baquerizo, M., Egidi, E., Guirado, E., Leach, J. E., Liu, H., & Trivedi, P. (2023). Climate change impacts on plant pathogens, food security and paths forward. Nature Reviews Microbiology 21:640–656.

Sapkota, R., Knorr, K., Jørgensen, L. N., O’Hanlon, K. A., & Nicolaisen, M. (2015). Host genotype is an important determinant of the cereal phyllosphere mycobiome. New Phytologist 207:1134–1144.

Tedersoo, L., Sánchez-Ramírez, S., Kõljalg, U., Bahram, M., Döring, M., Schigel, D., May, T., Ryberg, M., & Abarenkov, K. (2018). High-level classification of the Fungi and a tool for evolutionary ecological analyses. Fungal Diversity 90:135–159.

Testempasis, S. I., Papazlatani, C. V., Theocharis, S., Karas, P. A., Koundouras, S., Karpouzas, D. G., & Karaoglanidis, G. S. (2023) Vineyard practices reduce the incidence of *Aspergillus* spp. and alter the composition of carposphere microbiome in grapes (*Vitis vinifera* L.). Frontiers in Microbiology 14:1257644.

Tiedje, J. M., Bruns, M. A., Casadevall, A., Criddle, C. S., Eloe-Fadrosh, E., Karl, D. M., Nguyen, N. K., & Zhou J. 2022. Microbes and climate change: a research prospectus for the future. mBio 13:e00800–22.

Toju, H., Tanabe, A. S., Yamamoto, S., & Sato, H. (2012). High-coverage ITS primers for the DNA-based identification of ascomycetes and basidiomycetes in environmental samples. PLoS One 7:e40863.

van Leeuwen, C., & Seguin, G. (2006). The concept of terroir in viticulture. Journal of Wine Research 17:1–10.

Vasseur, L., & Catto, N. (2007). Atlantic Canada. In Lemmen, D. S., Warren, F. J., & Lacroix, J. (Eds.), From Impacts to Adaptation: Canada in a Changing Climate. Government of Canada, Ottawa, ON.

Weiss, M., Bauer, R., Sampaio, J. P., & Oberwinkler, F. (2014). *Tremellomycetes* and related groups. In: McLaughlin, D., Spatafora, J. (eds) Systematics and Evolution. The Mycota, vol 7A. Springer, Berlin, Heidelberg.

Wicaksono, W. A., Morauf, C., Müller, H., Abdelfattah, A., Donat, C., & Berg, G. (2023) The mature phyllosphere microbiome of grapevine is associated with resistance against *Plasmopara viticola*. Frontiers in Microbiology 14:1149307.

Wickham, H. (2016). Ggplot2: elegant graphics for data analysis. Springer-Verlag New York.

Wilcox, W. F., Gubler, W. D., & Uyemoto J. K. (2015) Compendium of Grape Diseases, Disorders, and Pests. (2^nd^ ed.) The American Phytopathological Society Press.

Zhang, H., Godana, E. A., Sui, Y., Yang, Q., Zhang, X., & Zhao L. (2020) Biological control as an alternative to synthetic fungicides for the management of grey and blue mould diseases of table grapes: a review. Critical Reviews in Microbiology 46:450–462.

Zhou, J., Cavagnaro, T. R., De Bei, R., Nelson, T. M., Stephen, J. R., Metcalfe, A., Gilliham, M., Breen, J., Collins, C., & López, C. M. R. (2021) Wine terroir and the soil bacteria: an amplicon sequencing–based assessment of the Barossa Valley and its sub-regions. Frontiers in Microbiology 11:597944.

Zhu, Y. G., Xiong, C., Wei, Z., Chen, Q. L., Ma, B., Zhou, S. Y. D., Tan, J., Zhang, L. M., Cui, H. L., & Duan, G. L. (2022) Impacts of global change on the phyllosphere microbiome. New Phytologist 234:1977–1986.

