## Supplementary Materials for "Experimental field trials model how the climate crisis will alter the phyllosphere and carposphere fungal communities of *Vitis* sp. L’Acadie Blanc"

### **Supplementary Data**

DADA2 Workflow

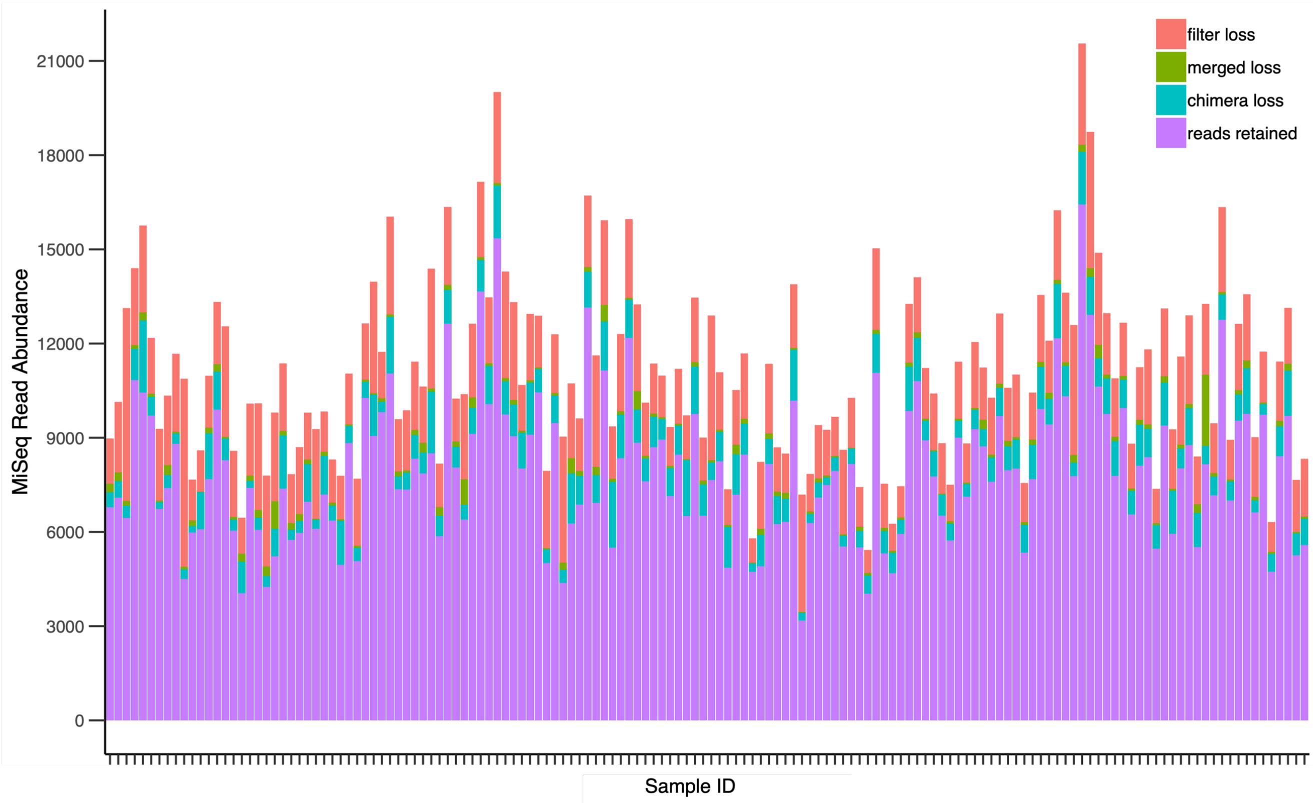


**Figure S1**. The DADA2 workflow processed 1 616 465 raw ITS reads from *Vitis* sp. grown during the 2020 season in Wolfville, Nova Scotia, produced from one lane of sequencing via Illumina's MiSeq at Génome Québec, and retained 1 160 088 reads which were used to infer amplicon sequence variants (ASVs).

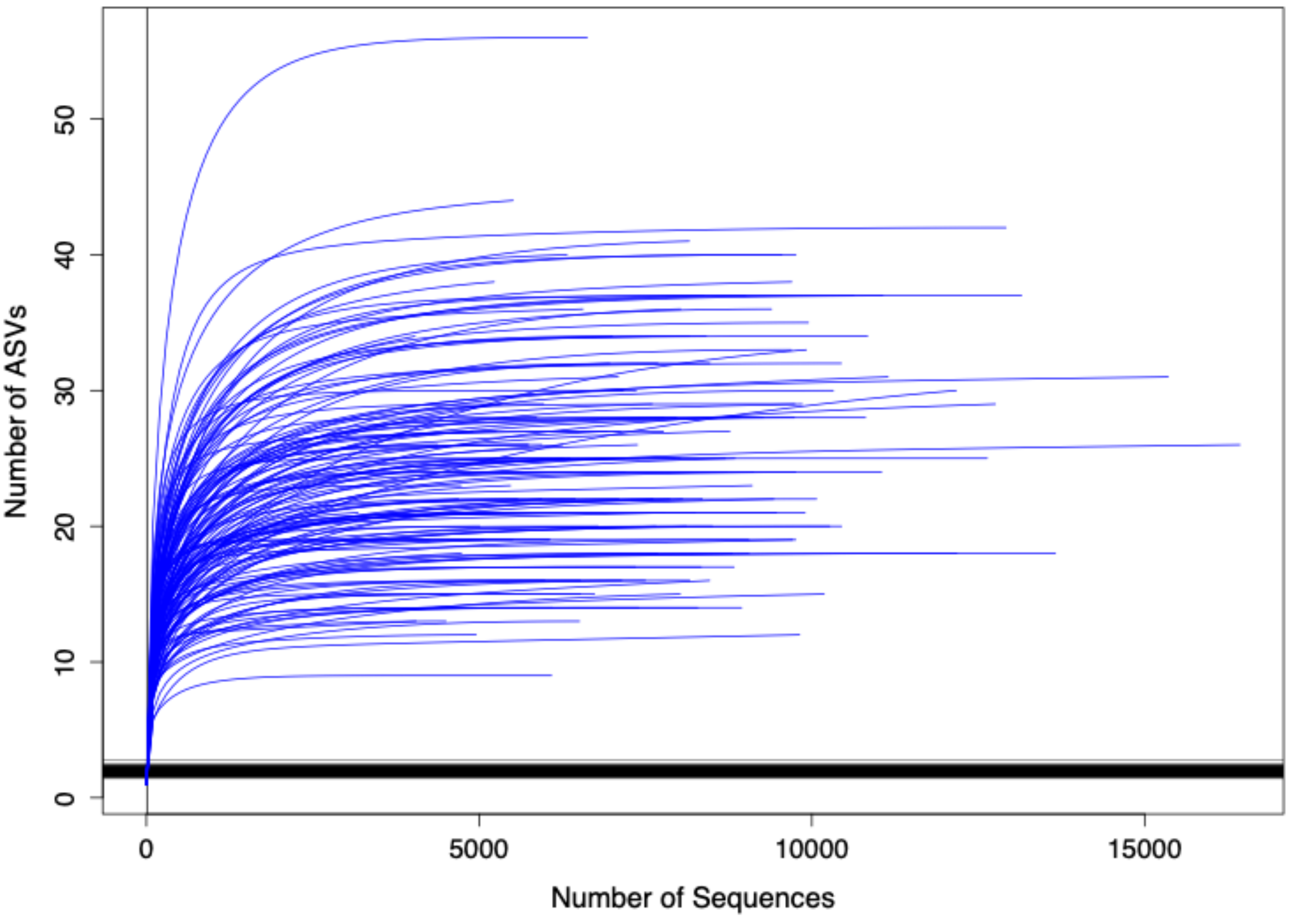


**Figure S2**. Rarefaction curves illustrated that the majority of the fungal communities were identified from our Illumina MiSeq sequencing depth in the samples harvested from the leaves and fruits of *Vitis* sp. grown during the 2020 season in Wolfville, Nova Scotia.

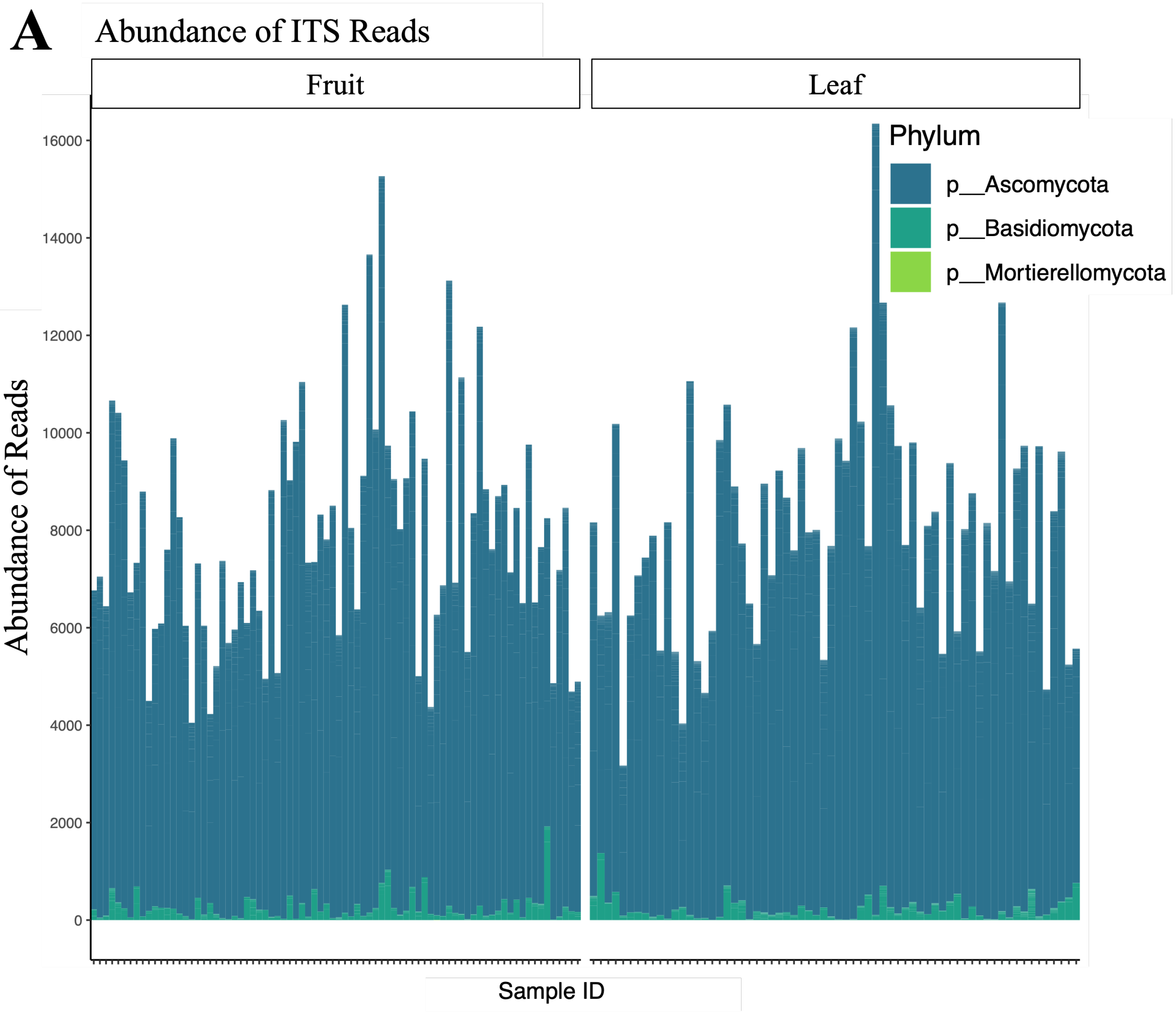
A
Abundance of ITS Reads
Fruit
Leaf
Abundance of Reads
16000
14000
12000
10000
8000
6000
4000
2000
0
Sample ID
Phylum
p__Ascomycota
p__Basidiomycota
p__Mortierellomycota

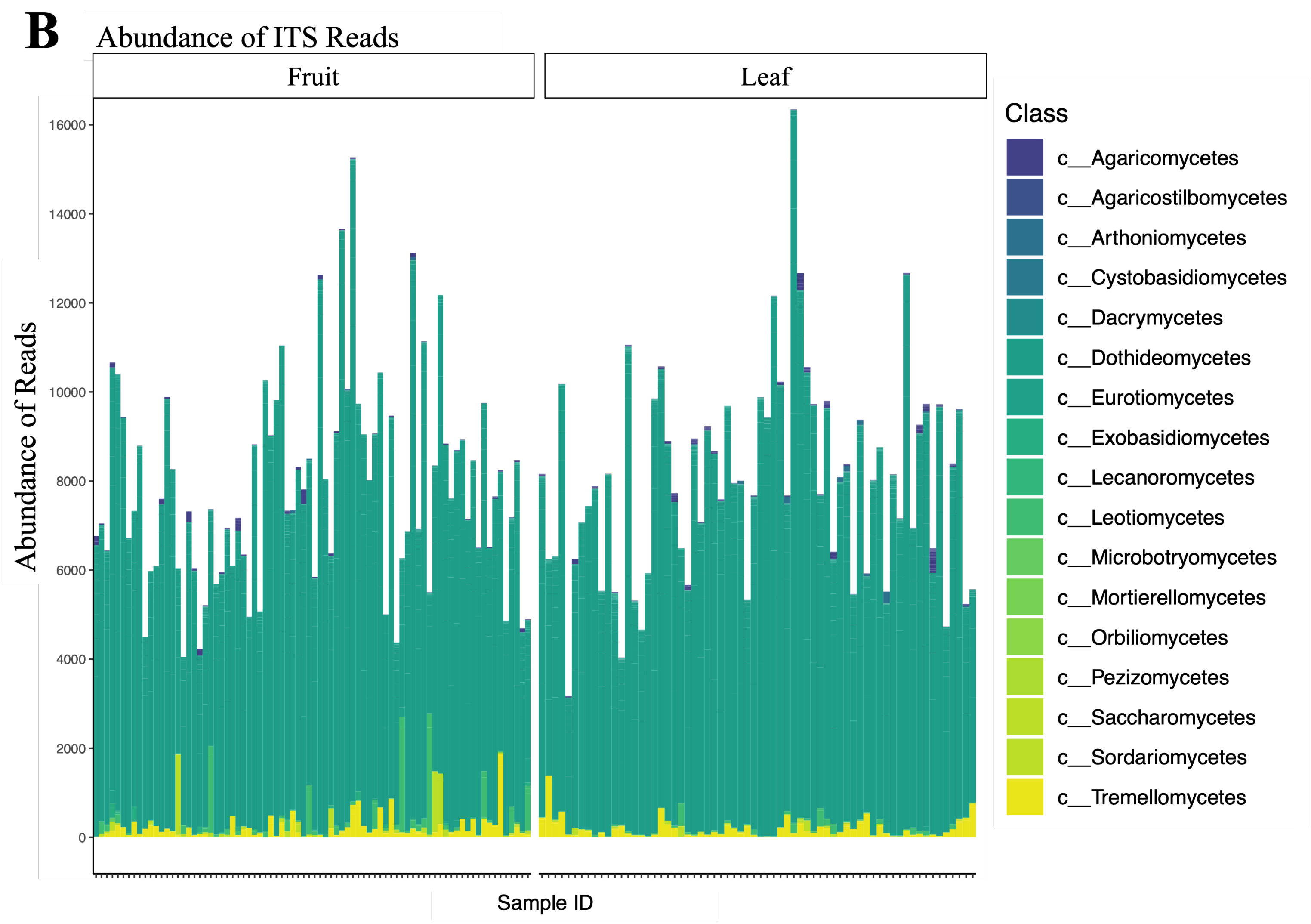
B
Abundance of ITS Reads
Fruit
Leaf
Abundance of Reads
16000
14000
12000
10000
8000
6000
4000
2000
0
Sample ID
Class
c__Agaricomycetes
c__Agaricostilbomycetes
c__Arthoniomycetes
c__Cystobasidiomycetes
c__Dacrymycetes
c__Dothideomycetes
c__Eurotiomycetes
c__Exobasidiomycetes
c__Lecanoromycetes
c__Leotiomycetes
c__Microbotryomycetes
c__Mortierellomycetes
c__Orbiliomycetes
c__Pezizomycetes
c__Saccharomycetes
c__Sordariomycetes
c__Tremellomycetes

**Figure S3**. Fungal community composition, represented here as phyla (A) and class (B) from the leaves and fruits of *Vitis* sp. cv. L'Acadie blanc sampled across the 2020 season in Wolfville, Nova Scotia. High-quality MiSeq reads from the ITS amplicons were retained through the DADA2 pipeline, inferred as amplicon sequence variants (ASVs), and assigned taxonomy using the UNITE database. The total dataset shows that the fungal leaf and fruit communities were dominated by Ascomycetes (A), of the class *Dothideomycetes* (B).

Differential Abundance Between Fungal Communities ($p <$ 0.05)

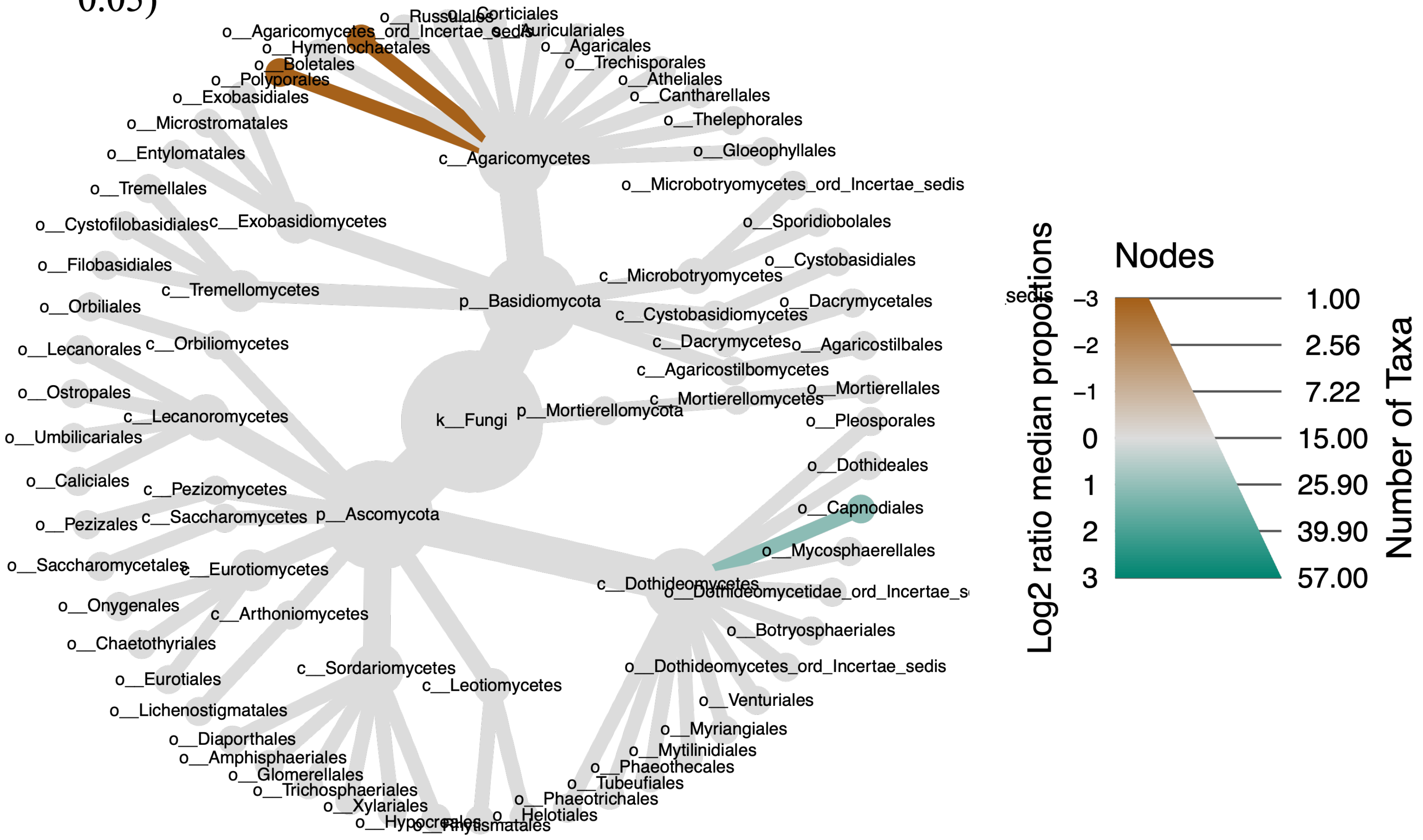

**Figure S4**. The fungal communities of the *Vitis* sp. leaves (green) were enriched in taxa from the *Capnodiales* order, while the fruit (brown) communities were enriched in *Polysporales* and *Agaricomycetes*. Samples were grown and harvested during the 2020 season in Wolfville, Nova Scotia. cladosporiaceae
